# Single-soma transcriptomics of tangle-bearing neurons in Alzheimer’s disease reveals the signatures of tau-associated synaptic dysfunction

**DOI:** 10.1101/2020.05.11.088591

**Authors:** Marcos Otero-Garcia, Yue-Qiang Xue, Tamara Shakouri, Yongning Deng, Samuel Morabito, Thomas Allison, William E. Lowry, Riki Kawaguchi, Vivek Swarup, Inma Cobos

## Abstract

Aggregation of hyperphosphorylated tau in neurofibrillary tangles (NFTs) is closely associated with neuronal death and cognitive decline in Alzheimer’s disease (AD). To define the signatures that distinguish between aggregation-prone and resistant cell states in AD, we developed a FACS-based method for the high-throughput isolation and transcriptome profiling of individual cells with cytoplasmic aggregates and profiled 63,110 somas from human AD brains. By comparing NFT-bearing and NFT-free somas within and across neuronal subtypes, we identified the cell-type-specific and shared states. NFT-bearing neurons shared a marked upregulation of genes associated with synaptic transmission, including a core set of 63 genes enriched for synaptic vesicle cycle and transsynaptic signaling, whereas glucose metabolism and oxidative phosphorylation changes were highly neuronal-subtype-specific. Apoptosis was modestly enriched in NFT-bearing neurons despite the strong link between tau and cell death. Our datasets provide a resource for investigating tau-mediated neurodegeneration and a platform for biomarker and drug target discovery.

## Introduction

Alzheimer’s disease (AD) pathology is defined by the combined presence of extracellular amyloid-β (Aβ) plaques and intracellular hyperphosphorylated tau aggregates in neurofibrillary tangles (NFTs) (Braak and Braak, 1991; Hyman et al., 2012; Serrano-Pozo et al., 2011). In the neocortex, amyloid plaque buildup begins years before the onset of cognitive deficits (Braak and Del Tredici, 2015; Nelson et al., 2012), whereas NFTs appear later and progress in close association with cognitive decline, loss of synapses, and neuronal cell death (Ballatore et al., 2007; Jack et al., 2010; Nelson et al., 2012; Terry et al., 1991). Tau hyperphosphorylation appears to contribute to disease pathogenesis by disrupting axonal transport and synapse function, affecting the cellular stress response, and promoting neuroinflammation (Busche et al., 2019; Sherman et al., 2016; Zhou et al., 2017). Notably, pathological tau has been shown to spread through synapses and functionally connected brain regions, contributing to disease progression (Franzmeier et al., 2019; Gibbons et al., 2019). The precise identity of the neurons that develop tau pathology in human brain and their molecular signatures, however, remain poorly characterized. Here, we developed a FACS-based method for the high-throughput isolation of individual somas with NFTs from postmortem human AD brain and applied single-soma transcriptomics to identify the precise excitatory and inhibitory neuronal subpopulations that exhibit NFTs and to characterize the molecular signatures of NFT susceptibility within and across neuronal subtypes.

Previous studies identified the brain regions and some cell types that are vulnerable to tau pathology in AD, including entorhinal cortex layer 2 pyramidal neurons and hippocampal CA1 pyramidal neurons (Gómez-Isla et al., 1996). In the neocortex, the vulnerability of specific neuronal subtypes is suggested by the laminar distribution of NFTs in specific layers, particularly in layers 2-3 and 5 (Bussière et al., 2003; Hof et al., 1990). Quantitative immunohistochemical and *in situ* hybridization (ISH) studies demonstrated the vulnerability of excitatory projection neurons to tau pathology compared to GABAergic inhibitory interneurons (Fu et al., 2019; Hof et al., 1991; Hof et al., 1993; Saiz-Sanchez et al., 2015). Although bulk RNA-sequencing (RNA-seq) and network-based analysis have implicated microtubule-regulating pathways (Roussarie et al., 2018), heat shock response and autophagy (Fu et al., 2019) in the vulnerability to tau, they have not resolved the heterogeneity of cellular and transcriptional responses associated with tau pathology in human AD.

Transcriptome profiling of single cells or single nuclei is a powerful approach to resolve cellular heterogeneity and define pathological cellular states (Darmanis et al., 2015; Habib et al., 2017; Hodge et al., 2019; Lake et al., 2016). Notably, studies directly comparing gene expression data from nuclei versus whole cells have shown a high degree of concordance (Bakken et al., 2018; Grindberg et al., 2013; Lake et al., 2017). Single-nucleus RNA-seq has been successfully applied to frozen human AD brains, revealing shared and cell-type-specific gene expression changes, sex-biased transcriptional responses, and potential drivers of disease progression (Del-Aguila et al., 2019; Grubman et al., 2019; Mathys et al., 2019). Nuclear profiling, however, cannot distinguish cells with and without cytoplasmic aggregates such as NFTs. Profiling whole cells from fresh brain tissue is feasible (Darmanis et al., 2015); however, single-cell suspensions from fresh brain are typically low yield and biased towards a recovery of glial cells, the harsh enzymatic digestion required for cell dissociation introduces transcriptome alterations, and obtaining fresh brain tissue is a major limitation for studying archived material and uncommon clinical samples.

Here, we developed new procedures for the high-throughput isolation of individual somas with NFTs from frozen human brains and profiled the transcriptomes of 63,110 somas with or without NFTs from the prefrontal cortex of Braak VI AD donors. By comparing the transcriptomes of single neurons with NFTs to those of neighboring NFT-free neurons, we provide unbiased and precise identification of the neuronal subtypes exhibiting aggregates. Differential gene expression (DGE) analysis between somas with or without NFTs within each neuronal subtype and across subtypes identified the cell-type-specific and commonly altered genes and pathways associated with pathological tau. NFT-bearing neurons across subtypes shared a marked upregulation of genes associated with synaptic transmission, particularly the synaptic vesicle cycle and transsynaptic signaling. We report a ranked list of 227 synaptic genes dysregulated in NFT-bearing neurons, including a core set of 63 genes shared across subtypes. The well-known biomarkers of neurodegeneration and cognitive decline NEFL, SNAP25 and SYT1 (Brinkmalm et al., 2014; Davidsson et al., 1996; Galasko et al., 2019; Mattsson et al., 2017; Mattsson et al., 2019; Preische et al., 2019) ranked within the top 25, highlighting the value of our datasets for discovery. Our data provide a framework that can be applied to disease modeling and the discovery of new biomarkers and drug targets.

## Results

### Isolation and transcriptome profiling of single somas with NFTs from human AD brains

Because the standard methods for profiling single nuclei from the postmortem human brain (Habib et al., 2017; Krishnaswami et al., 2016; Lake et al., 2016; Mathys et al., 2019) cannot distinguish between cells with and without cytoplasmic aggregates, we developed procedures for the high-throughput isolation of NFT-bearing somas from fresh-frozen tissue (Figure 1A). We focused on the prefrontal cortex (Brodmann area 9; BA9) of Braak stage VI AD patients, where NFT-bearing neurons represent 5–8% of the total neuronal population (Figure 1B). We microdissected the cortical ribbon spanning all cortical layers and performed gentle mechanical dissociation, without the use of detergents or enzymatic digestion, followed by sucrose-iodixanol gradient centrifugation (Figure 1C). Immunostaining and fluorescence-activated cell sorting (FACS) using the AT8 antibody, which detects hyperphosphorylated tau aggregates, and the pan-neuronal marker MAP2 allowed us to isolate single NFT-bearing somas (AT8^+^) and neighboring NFT-free somas (MAP2^+^/AT8^−^; referred to as MAP2^+^) from the same homogenate (Figure 1B). Microscopic examination of the sorted AT8^+^ population confirmed the efficient isolation of somas with NFTs (Figure 1C).

**Figure 1.**
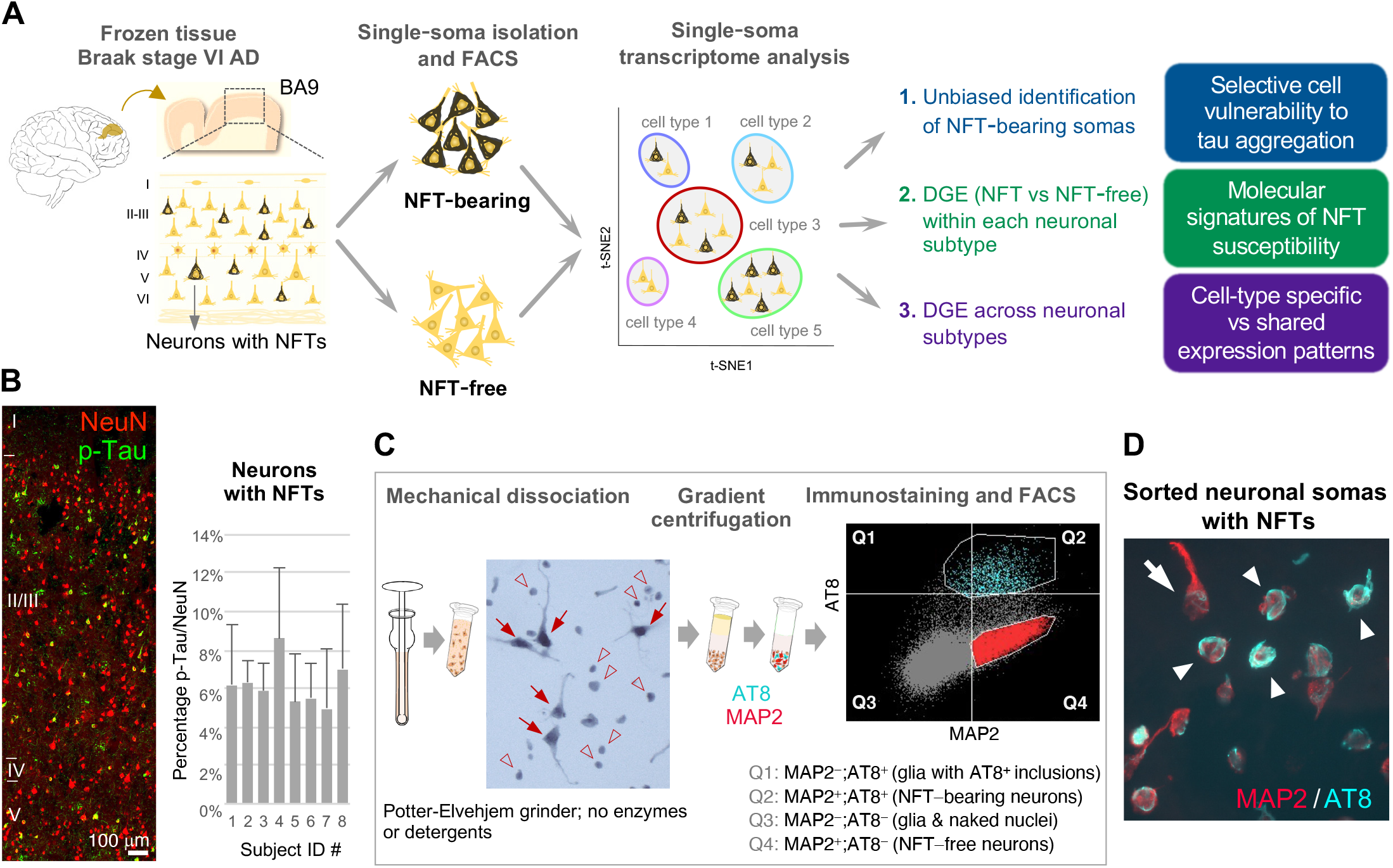
Isolation and transcriptome profiling of single somas with NFTs from human AD brains. (A) Overview of our strategy to compare the transcriptomes of somas with and without NFTs from the human AD brain. (B) Quantification of NFT-bearing neurons in the Braak VI AD prefrontal cortex (BA9). A representative section immunostained with NeuN and p-Tau illustrates neurons with NFTs predominantly in layers 2-3 and 5. The bar chart shows the percentages of neurons with NFTs in eight donors (9–11 counting frames per case; 2,000–3,000 NeuN^+^ cells; ~4.5 mm^2^ total area); error bars indicate standard deviation. The mean proportion of neurons with NFTs and standard deviation was 6.3 ± 1.15%. (C) Experimental approach to single-soma isolation. Gentle mechanical force without detergents or enzymatic digestion released cells with relatively well preserved somas (arrows) and naked nuclei (arrowheads). After gradient centrifugation, somas were stained with SYTOX green nucleic acid stain and MAP2 and AT8 antibodies. FACS gates were set to collect all cells with tau aggregates (AT8^+^; blue subpopulation) and neurons without aggregates (MAP2^+^/AT8^−^; red subpopulation). (D) Representative sorted AT8^+^/MAP2^+^ neurons displaying characteristic NFTs, including mature tangles (band or flamed shaped; arrowheads) and early perikaryal fibrils (arrow). See also Figure S1.

To compare our method for single-soma isolation with standard single-nucleus isolation, we processed single-nucleus and single-soma suspensions prepared from the same frozen tissue sample side by side. We used the pan-neuronal markers NeuN (nuclear) and MAP2 (cytoplasmic) to sort by FACS single nuclei or somas, respectively, and analyzed the transcriptomes of 13,194 cells (Figure S2A). MAP2^+^ cells showed modest increases in the median number of unique molecular identifiers (UMIs) per cell (1,501 vs. 1,259 for NeuN), the percentage of exonic reads (46.7% vs. 42.4% for NeuN), and mitochondrial gene content (14.9% vs. 10.8% for NeuN) (Figures S2B–D). These metrics indicate efficient profiling of nuclear transcripts using both methods and a limited contribution of cytoplasmic transcripts from MAP2^+^ cells. This outcome is expected for frozen tissues in which cell membranes have been disrupted and thus cytoplasmic transcripts have been largely lost. Notably, the ability to discriminate neuronal subtypes was highly similar (Figures S2E–H). Thus, our method allows the comparison of somas with and without NFTs from the same tissue sample, which overcomes confounding effects caused by differences in genetic background, sex, age, comorbidities, medication use, premortem agonal state, and tissue processing procedures.

### Cell type composition of the NFT-free and NFT-bearing datasets from human AD brain

In the brains of AD patients, tau aggregates are found in neurons and only rarely in glia. In the cerebral cortex, in particular, NFTs develop in the somas of pyramidal excitatory neurons (Braak and Braak, 1991; Fu et al., 2019; Hof et al., 1991; Hof et al., 1993; Hyman et al., 2012; Saiz-Sanchez et al., 2015). The specific neuronal subtypes vulnerable to tau pathology are, however, not fully resolved. To determine the cellular specificity of pathological tau aggregates, we analyzed the transcriptomes of 24,660 NFT-bearing somas (AT8^+^) and 38,465 neighboring NFT-free somas (MAP2^+^) from the BA9 of eight Braak VI AD patients. All the patients suffered from dementia and received a neuropathological score of A3B3C3 (Hyman et al., 2012) at the time of death (Table S1). The samples were sequenced in a single batch at a depth of ~72,000 reads per soma, corresponding to a sequencing saturation of ~85%. The median numbers of genes and UMIs per soma were higher in the AT8^+^ somas (1,481 and 2,440, respectively) than in MAP2^+^ (1,248 and 2,062, respectively; Figures 2B and 2E and Table S2) somas. This increase in UMIs and genes per soma in the NFT dataset may have resulted from differences in cell composition (i.e, larger cells have higher transcript abundance) and/or transcriptional upregulation.

**Figure 2.**
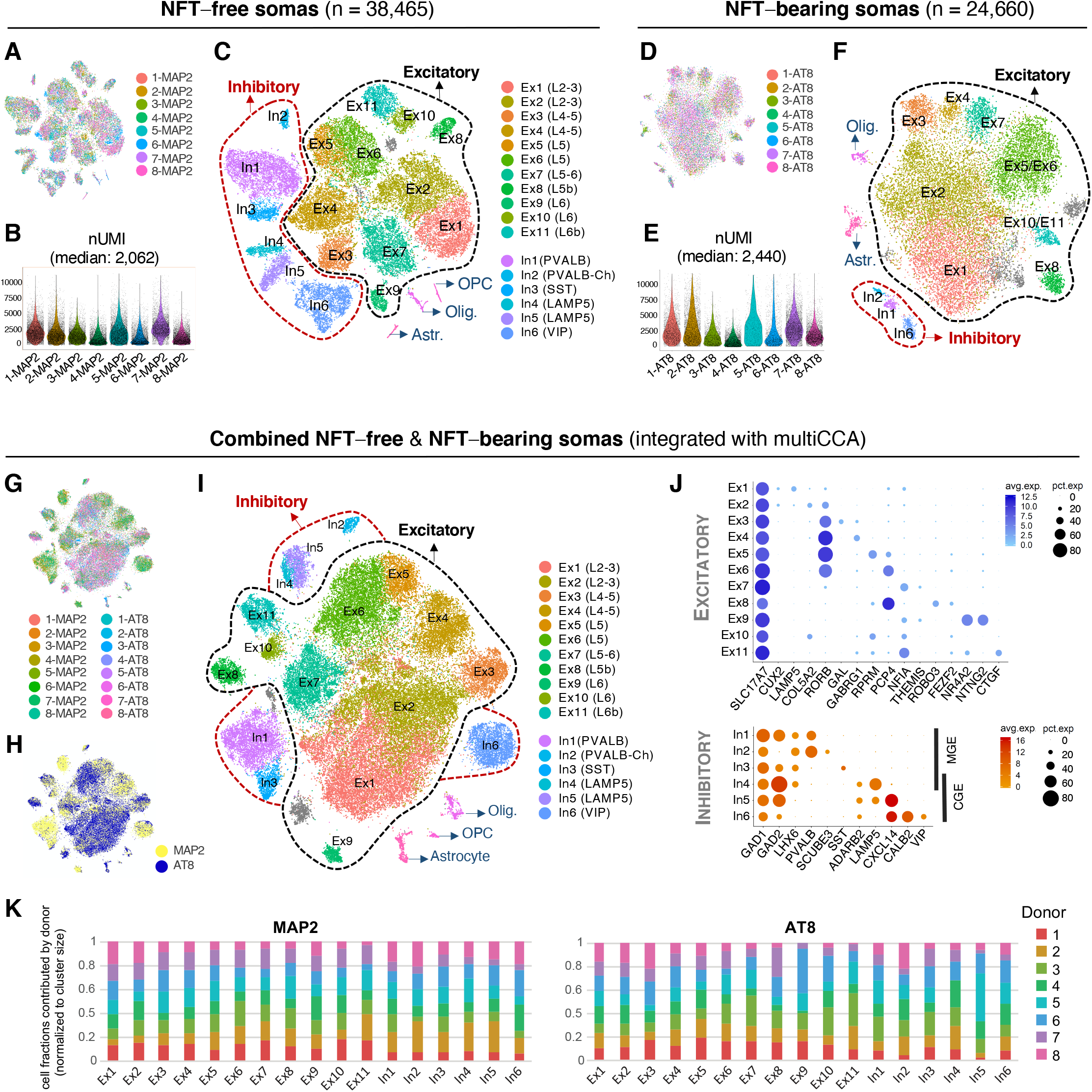
Cell type composition of the NFT-free and NFT-bearing datasets from human AD brain. (A–F) Profiling of NFT-free (MAP2; A–C) and NFT-bearing (AT8; D–F) somas isolated from the prefrontal cortex of eight Braak VI AD donors. t-SNE plots show unsupervised clustering of single somas. Violin plots show the distribution of UMI counts per cell in each sample. The NFT-free dataset (C) contained 11 excitatory neuron clusters (Ex1 to Ex11; 27,080 cells), 6 inhibitory neuron clusters (In1 to In6; 10,232 cells), and 3 small glial clusters (astrocytes, oligodendroglia, and OPCs; 206, 307, and 220 cells, respectively). The NFT dataset (F) contained primarily excitatory neurons (8 clusters; 21,567 cells) and small clusters of inhibitory neurons (739 cells) and glia (610 cells). The gray clusters represent mixed populations. (G–J), Analysis of the AT8 (NFT-bearing) and MAP2 (NFT-free) combined datasets (n = 63,110 somas) after multiCCA analysis. Colors in (G) correspond to each sample. The t-SNE plot in (H) highlights individual MAP2^+^ (yellow) and AT8^+^ (blue) somas. Unsupervised clustering (I) identified the same neuronal subtypes as in the NFT-free (MAP2-only) dataset (C). The dot plots (J) depict the expression of marker genes (x-axis) within excitatory (top) and inhibitory (bottom) neuron clusters (y-axis). Inhibitory neurons included LHX6^+^ and ADARB2^+^ subpopulations, which originate from the medial or caudal ganglionic eminence (MGE or CGE), respectively. The sizes of the dots represent the percentage of cells expressing the marker; color intensities represent the expression level. (K) Bar charts illustrating the fraction of somas derived from each donor (normalized to cluster size). Bars on the x-axis correspond to clusters (excitatory Ex1 to Ex11 and inhibitory In1 to In6). Colors correspond to each donor. See also Figures S2 and S3, and Tables S2 and S3.

To determine the cell composition of the NFT-free and NFT-bearing datasets, we used an unbiased approach. Unsupervised clustering of the NFT-free (MAP2^+^) somas identified a cell type composition that was concordant with that described in previously published datasets from control and AD nuclei (Hodge et al., 2019; Mathys et al., 2019) (Figures 2A–C and S3). Using conservative parameters, we annotated 17 clusters that corresponded to 11 distinct excitatory cell subtypes (referred to as Ex1 to Ex11) expressing the pan-excitatory marker SLC17A7, and 6 inhibitory cell subtypes (referred to as In1 to In6) expressing the pan-inhibitory marker GAD1 (Figure S4; Table S3). Excitatory subtypes included layers 2-3 CUX2^+^ (composed of superficial LAMP5^+^/SERPINE^+^ [Ex1] and deeper COL5A2^+^ cells [Ex2]); layers 4-5 RORB^+^ (RORB^+^/GAL^+^, RORB^+^/GABGR1^+^, RORB^+^/RPRM^+^, and RORB^+^/PCP4^+^ [Ex3 to E6; respectively]); layer 5b PCP4^+^/ROBO3^+^ [Ex8]; layers 5-6 NFIA^+^/THEMIS^+^ [Ex7]; layer 6 NR4A2^+^/NTNG2^+^ [Ex9] and FEZF2^+^/COL5A2^+^ [Ex10]; and the deeper layer 6b FEZF2^+^/CTGF^+^ [Ex11] cells. Inhibitory interneurons included two major classes with developmental origins in the medial (LHX6^+^) or caudal (ADARB2^+^) ganglionic eminences. LHX6^+^ cells consisted of PVALB^+^ and SST^+^ subtypes. A distinct PVALB^+^ cluster characterized by high expression of GAD1, low expression of GAD2 and expression of SCUBE3 defined putative chandelier cells (Hodge et al., 2019). ADARB2^+^ cells included LAMP5^+^/KIT^+^ cells (split into two clusters by the expression of CXCL14) and a highly heterogeneous VIP^+^/CALB2^+^ cluster. The proportion of inhibitory cells was 26.6%. Thus, all major excitatory and inhibitory neuronal subtypes were identified in prefrontal cortex samples from Braak VI AD patients, despite the advanced stage of neurodegeneration.

Unsupervised clustering of the NFT somas (AT8^+^) using the same analysis pipeline as for the NFT-free somas showed clusters corresponding to cell types and technical covariates, including clusters derived from individual donors. To improve data integration, we used multiple canonical correlation analysis (multiCCA) (Butler et al., 2018). This analysis removed the clusters originating from individual donors and enhanced cell identity-based clustering (Figures 2D–F). Although the FACS windows were set to capture both neuronal and nonneuronal cells, the latter consisted of a small fraction (2.7%), as expected due to the rare occurrence of tau aggregates in glia in human AD (Braak and Braak, 1991; Serrano-Pozo et al., 2011). We annotated 8 excitatory neuron clusters, 3 inhibitory neuron clusters, and 2 glial clusters (Figure 2F; Table S3). Approximately 94% of the AT8^+^ somas were excitatory neurons, and 3.2% were interneurons. The excitatory clusters in the NFT dataset were less distinct than those in the NFT-free dataset, likely resulting from cell state variation associated with NFT pathology superimposed on the variation driven by cell identity. Some excitatory subtypes were not identified, likely due to the small numbers of cells for that particular subtype.

We next analyzed the AT8^+^ and MAP2^+^ somas together using multiCCA (Figures 2G–J). Multidataset integration by multiCCA enhanced the identification of neuronal subtypes and reduced the effects of cell state variation. We identified the same 17 neuronal subtypes as in the MAP2^+^ dataset (Figures 2C and 2I). Clustering was not driven by particular samples, as each of the eight donors contributed cells evenly to the clusters (Figure 2K). This clustering approach allowed us to match the NFT-bearing and NFT-free somas of the same subtype for subsequent cell counting and DGE analysis.

### Census of neuronal subtypes exhibiting NFTs in human AD

To obtain an accurate census of the neuronal subtypes exhibiting NFTs, we counted the numbers of somas with and without NFTs within each cluster identified in the integrated AT8^+^ and MAP2^+^ dataset (Figures 3A–C; Table S3). The proportions of neurons with NFTs ranged from 1.0–11.7% for the excitatory subtypes and from 0.5–6.8% for the inhibitory subtypes (Figure 3B), after normalization to NFT densities in tissue (Figure 1B). The excitatory subtypes with the highest proportions of NTFs were the layers 2-3 Ex1 and Ex2 (11.7% and 10.5%, respectively) and the layer 5 Ex6 (10.6%) (Figure 3C). Notably, layer 5 was highly heterogenous and contained vulnerable (Ex6; 10.6%) and relatively resistant (Ex4 and Ex5; 3.6% and 5.2%, respectively) populations. Layer 6 was largely spared although a deep layer 6b subpopulation showed intermediate proportions of NFTs (Ex11; 5.6%). Most inhibitory neurons were spared (overall 1.9%), with the exception of chandelier cells (In2; 6.8%) (Figure 3C). Thus, different subpopulations or excitatory and inhibitory neurons demonstrate different susceptibilities to NFT formation. The results were consistent across clustering algorithms and robust to variation in the clustering parameters (Figure S4).

**Figure 3.**
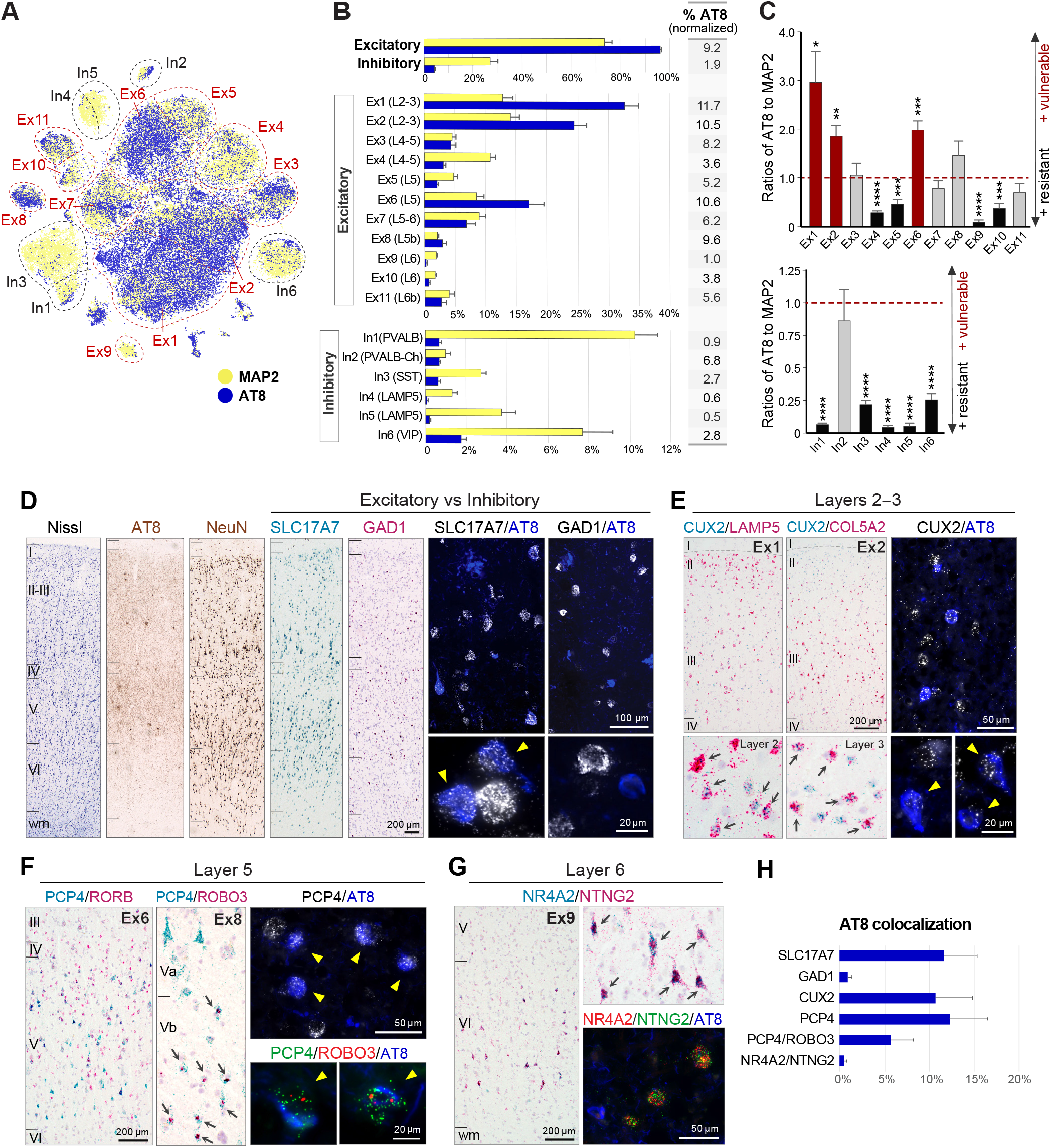
Census of neuronal subtypes exhibiting NFTs in human AD. (A) t-SNE plot highlighting the relative contributions of MAP2^+^ (yellow) and AT8^+^ (blue) somas to each cluster. (B) Bar plots showing the percentages of MAP2^+^ and AT8^+^ somas contributing to each cluster. Solid bars represent the average percentages of the eight samples. Error bars indicate standard error of the mean. The adjacent column shows the expected percentages of AT8^+^ somas in histological sections (data normalized to NFT densities from Figure 1B). (C) Bar plots showing the relative vulnerabilities to NFTs among excitatory and inhibitory clusters. Dashed lines indicate an AT8 to MAP2 ratio of 1:1. Red and black bars represent the most vulnerable and resistant neuronal subtypes, respectively (*p < 0.05, **p < 0.01, ***p < 0.001, and ****p < 0.0001 by one sample t-test against a hypothetical mean of 1). (D) Histological validation in Braak VI prefrontal cortex sections. Nissl, AT8, NeuN, and chromogenic ISH for marker genes provide anatomical reference and spatial resolution of gene expression. Dual fluorescent AT8 immunohistochemistry (blue) and ISH (white) stains illustrate the vulnerability of excitatory (SLC17A7^+^) relative to inhibitory (GAD1^+^) neurons. Yellow arrowheads point to individual neurons with NFTs. (E) Vulnerability of layers 2-3 CUX2^+^ excitatory neurons. Double chromogenic ISH for CUX2 (blue) and LAMP5 or COL5A2 (red) illustrate two distinct subpopulations with preferential distribution in superficial (CUX2^+^/LAMP5^+^; cluster Ex1) or deep (CUX2^+^/COL5A2^+^; cluster Ex2) layers. Arrows point to double-positive neurons. Dual fluorescent AT8 immunohistochemistry (blue) and CUX2 ISH (white) illustrate CUX2^+^ neurons with NFTs. (F) Vulnerability of layer 5 RORB^+^/PCP4^+^ (cluster Ex6) and layer 5b PCP4^+^/ROBO3^+^ (cluster Ex8) excitatory neurons. Double chromogenic ISH for PCP4 and ROBO3 illustrate a distinct subpopulation of small PCP4^+^/ROBO3^+^ neurons in layer 5b (arrows). Triple fluorescent AT8 immunohistochemistry (blue), PCP4 ISH (green), and ROBO3 ISH (red) illustrate neurons with NFTs. (G) Resistance of layer 6 NR4A2^+^/NTNG2^+^ (cluster Ex9) excitatory neurons. Double chromogenic and triple fluorescent stains illustrate NFT-free NR4A2^+^/NTNG2^+^ neurons. (H) Bar plot showing the frequencies of neurons with NFTs for each marker(s): SLC17A7^+^ (11.59 ± 3.7%); GAD1^+^ (0.95 ± 0.41%); CUX2^+^ (10.64 ± 4.2%); PCP4^+^ (12.24 ± 4.2%); PCP4^+^/ROBO3^+^ (5.70 ± 2.53%) and NR4A2^+^/NTNG2^+^ (0.5 ± 0.3%). Error bars indicate standard deviation. See also Figure S4.

We validated these findings using histological sections obtained from the same brains and tissue blocks (Figures 3D–H). Combined immunohistochemistry for AT8 and ISH for the markers SLC17A7 and GAD1 showed proportions of NFTs in excitatory neurons (11.59 ± 3.7%) and inhibitory neurons (0.95 ± 0.41%) similar to those in our transcriptome analysis (Figures 3D and 3H). Layers 2-3 excitatory neurons expressing the transcription factor CUX2, which coexpress either LAMP5 (Ex1) or COL5A2 (Ex2), showed high proportions of neurons with NFTs (10.64 ± 4.2%; Figures 3E and 3H). Layer 5 PCP4^+^ (Ex6) and layer 5b PCP4^+^/ROBO3^+^ (Ex8) subtypes were also vulnerable (12.24 ± 4.2% and 5.70 ± 2.53%, respectively; Figures 3F and 3H). In contrast, the layer 6 NR4A2^+^/NTNG2^+^ (Ex9) subtype was largely spared (0.5 ± 0.3%; Figures 3G and 3H). Collectively, these results indicate that tau pathology in AD shows marked neuronal subtype specificity at late stages of neurodegeneration and that this specificity can be effectively resolved by single-soma transcriptomics.

### Transcriptome signatures of tau pathology within and across excitatory neuronal subtypes

To characterize the transcriptomic changes associated with tau pathology, we performed a two-step DGE analysis, first between NFT-bearing and NFT-free within each neuronal subtype to define the molecular signatures of NFT susceptibility, and next comparing the lists of differentially expressed (DE) genes across subtypes to distinguish the shared and cell-type-specific transcriptomic changes. Using the MAST (model-based analysis of single-cell transcriptomics) generalized linear model (Finak et al., 2015) for DGE analysis within each subtype, we identified 692–978 DE genes in the clusters with high cell numbers and NFT proportions (Ex1, Ex2, Ex3, Ex6, Ex7) and 55–186 DE genes in the smaller clusters (Ex4, Ex5, Ex8, Ex11) (Figures 4A and 4B; Table S4) (MAST test; adjusted p-value < 0.05; log fold change > 0.1; detection in ≥ 20% of cells). Most of the DE genes were upregulated in the NFT neurons (~66–84% upregulated in Ex1, Ex2, Ex6, Ex7), except in cluster Ex3 (~39% upregulated). The widespread transcriptional upregulation is consistent with previous work showing a role for phosphorylated tau in chromatin remodeling (Frost et al., 2014; Klein et al., 2019). The overall transcriptomic changes associated with NFTs were robust to the subsampling of a subset of donors as assessed by the rank-rank hypergeometric overlap (RRHO) test (Figure S5).

**Figure 4.**
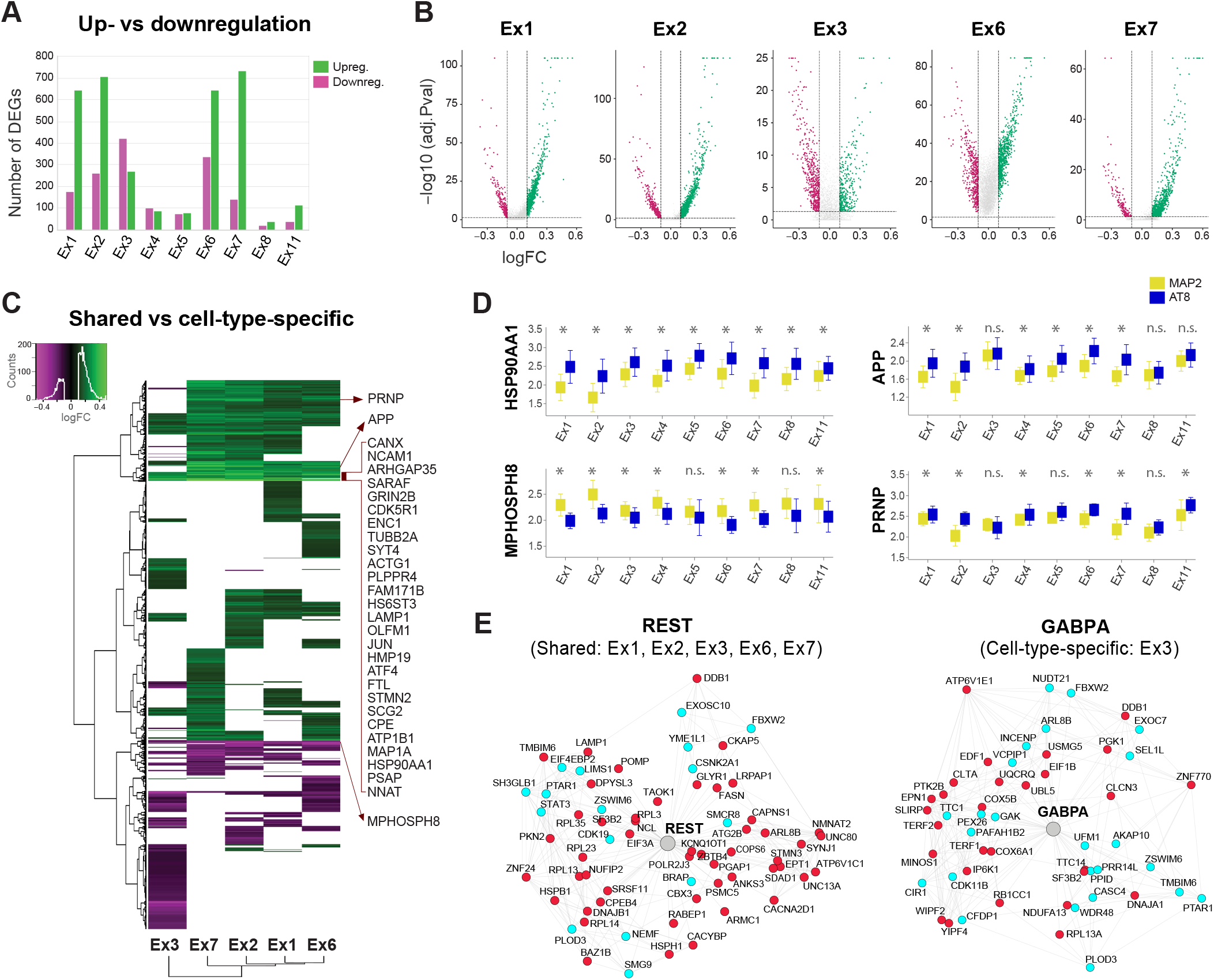
Transcriptome signatures of tau pathology within and across excitatory neuronal subtypes. (A) Total numbers of DE genes (green = upregulated; magenta = downregulated) between neurons with and without NFTs in nine excitatory neuronal subtypes (MAST test). (B) Volcano plots of the DE genes for each excitatory neuronal subtype. The y-axis corresponds to the adjusted p-values (capped to the 98^th^ percentile in each plot), and the x-axis corresponds to the log fold change. Dots represent genes (green = upregulated; magenta = downregulated; gray = not differentially expressed). Dashed lines indicate significance thresholds (log fold change values of < 0.1 or > 0.1; adjusted p-value < 0.05; for genes detected in ≥ 20% of cells in at least one condition). (C) Heatmap and hierarchical clustering of DE genes showing shared and cell type-specific upregulated (green) and downregulated (magenta) genes among five neuronal subtypes with high numbers of NFTs. Columns represent cell types; rows represent genes. The input lists of DE genes were truncated to match the cell type with the smallest number of DE genes (n = 692) to represent equal contributions from each of the five clusters. Of these 692 DE genes, 102 genes were upregulated in the five clusters; an additional 163 genes, including APP and PRNP, were upregulated in all five clusters except Ex3. A total of 22 genes were downregulated in the five clusters, including the epigenetic repressor MPHOSPH8. The genes listed by the heatmap include shared DE genes with high log fold change values. (D) Box plots showing the median expression values of HSP90AA1, MPHOSPH8, APP, and PRNP in neurons with and without NFTs in each excitatory cluster. Error bars indicate standard deviation. *p < 0.05 (MAST test); n.s. = not significant. (E) REST and GABPA transcriptional regulatory networks (color code: cyan = genes coregulated and coexpressed based on our single-cell data and/or ROSMAP datasets; red = genes coregulated, coexpressed, and differentially expressed in neurons with NFTs). See also Figures S5 and S6, and Tables S4 and S5.

We then performed hierarchical clustering of the DE genes across neuronal subtypes to distinguish the cell-type-specific and shared transcriptomic changes (Figure 4C; Table S5). We focused our analysis on the five excitatory clusters with the highest cell numbers and NFT proportions (E1, Ex2, Ex3, Ex6, Ex7) to avoid underestimating the extent of the shared transcriptomic changes (additional clusters are shown in an interactive hierarchical heatmap in Table S5). While many gene expression changes were neuronal subtype specific, this approach distinguished a subset of commonly altered genes in neurons with NFTs. A total of 124 DE genes (102 upregulated; 22 downregulated) were shared among all five clusters, and an additional 163 genes were upregulated in all clusters except Ex3 (Figure 4C). Among the commonly upregulated genes with the highest fold change were genes encoding synaptic proteins (i.e., CALM1, ATP1B1, GRIN2B, CDK5R1, SYT4, CANX, RTN4) and cytoskeletal proteins and microtubule dynamics regulators (i.e., ACTG1, TUBB2A, PLPPR4, MAP1A, ENC1, STMN2). Also commonly upregulated was NEFL, the gene encoding neurofilament light chain protein (NfL), a cerebrospinal fluid and plasma biomarker of neurodegeneration in AD patients (Mattsson et al., 2019). Other commonly upregulated genes included: the immediate early gene JUN; genes related to the integrated stress response, such as activating transcription factor 4 (ATF4); the gene encoding the heat shock protein and chaperone Hsp90 (HSP90AA1), which has been implicated in the intracellular processing of aggregated tau (Dickey et al., 2007) (Figure 4C); the gene encoding prosaposin (PSAP), a lysosomal protein and binding partner of progranulin (Nicholson et al., 2016); and the iron homeostasis-associated genes FTL and FTH1. Notably, APP, encoding the Aβ precursor protein, was upregulated in neurons with NFTs in most clusters but not in Ex3 (Figure 4D). Similar changes were observed for the prion protein-coding gene (PRNP; Figure 4D), which has been shown to act as a receptor for soluble Aβ oligomers (Laurén et al., 2009).

To investigate the gene regulatory networks underlying shared and cell-type-specific responses associated with pathological tau, we performed a transcription factor (TF) binding site enrichment analysis and generated TF regulatory networks integrating neuronal-specific ENCODE ChIP-seq data and consensus coexpression networks in AD (Mostafavi et al., 2018). Among the TF networks shared across neuronal subtypes was one centered around REST, a key regulator of neuronal differentiation and neuronal excitability that has been implicated in aging and AD (Lu et al., 2014; Zullo et al., 2019). Cluster Ex3 had a unique set of TF networks that included GABPA, a transcription factor involved in the regulation of mitochondrial genes required for electron transport and oxidative phosphorylation (Figure 4E and S6) (Yang et al., 2014). Together, our DGE and gene regulatory network analyses identified the shared and cell-type-specific molecular signatures associated with tau pathology across excitatory neuron subtypes.

### Shared versus cell-type-specific pathways associated with tau pathology

To analyze and visualize the shared and cell-type-specific pathways associated with tau pathology, we used functional enrichment analysis tools (Reimand et al., 2019) (Figure 5A). For this, we first generated a ranked list of statistically significant gene ontology (GO) terms enriched in neurons with NFTs for each cell type using g:Profiler (Figure 5B; Table S6), and then integrated the results from the five excitatory clusters with the highest cell numbers and NFT proportions (Ex1, Ex2, Ex3, Ex6, Ex7) into a single network using Cytoscape with EnrichmentMap. The resulting pathway enrichment map illustrates the pathways enriched in NFT neurons that are cell type specific or shared among cell types (Figure 5C) and delineates the set of genes associated with each dysregulated pathway (Table S7). The shared pathways across cell types with the highest enrichment scores were related to synaptic transmission (Figure 5C; Table S7). Other commonly enriched pathways in neurons with NFTs included calcium homeostasis, microtubule polymerization, axonal remodeling, dendritic spine remodeling, microtubule-based transport, and intracellular protein transport. In contrast, glucose metabolism and oxidative phosphorylation were highly cell type dependent and particularly enriched in the Ex3 cluster. Notably, neuronal cell death and apoptosis pathways were shared across cell types but represented modestly. Both pro-apoptotic and negative regulators of apoptosis pathways were represented. Genes in this category included FAIM2 and MIF (downregulated), and ATF4, BAD, BNIP3, and HIF1A (upregulated). A smaller set of genes involved in the regulation of mitochondrial membrane permeability transition associated with cell death were upregulated, including BAD, BNIP3, HSPA1A, and genes encoding 14-3-3 phospho-serine/phospho-threonine binding proteins (YWHAE, YWHAH, YWHAG, YWHAZ, YWHAB) (Figure 5C; Table S7). Collectively, our analysis highlights the shared and cell-type-specific pathways associated with tau pathology and reveals dysregulated pathways converging on the synapse in neurons with NFTs, supporting the view of AD as a synaptopathy.

**Figure 5.**
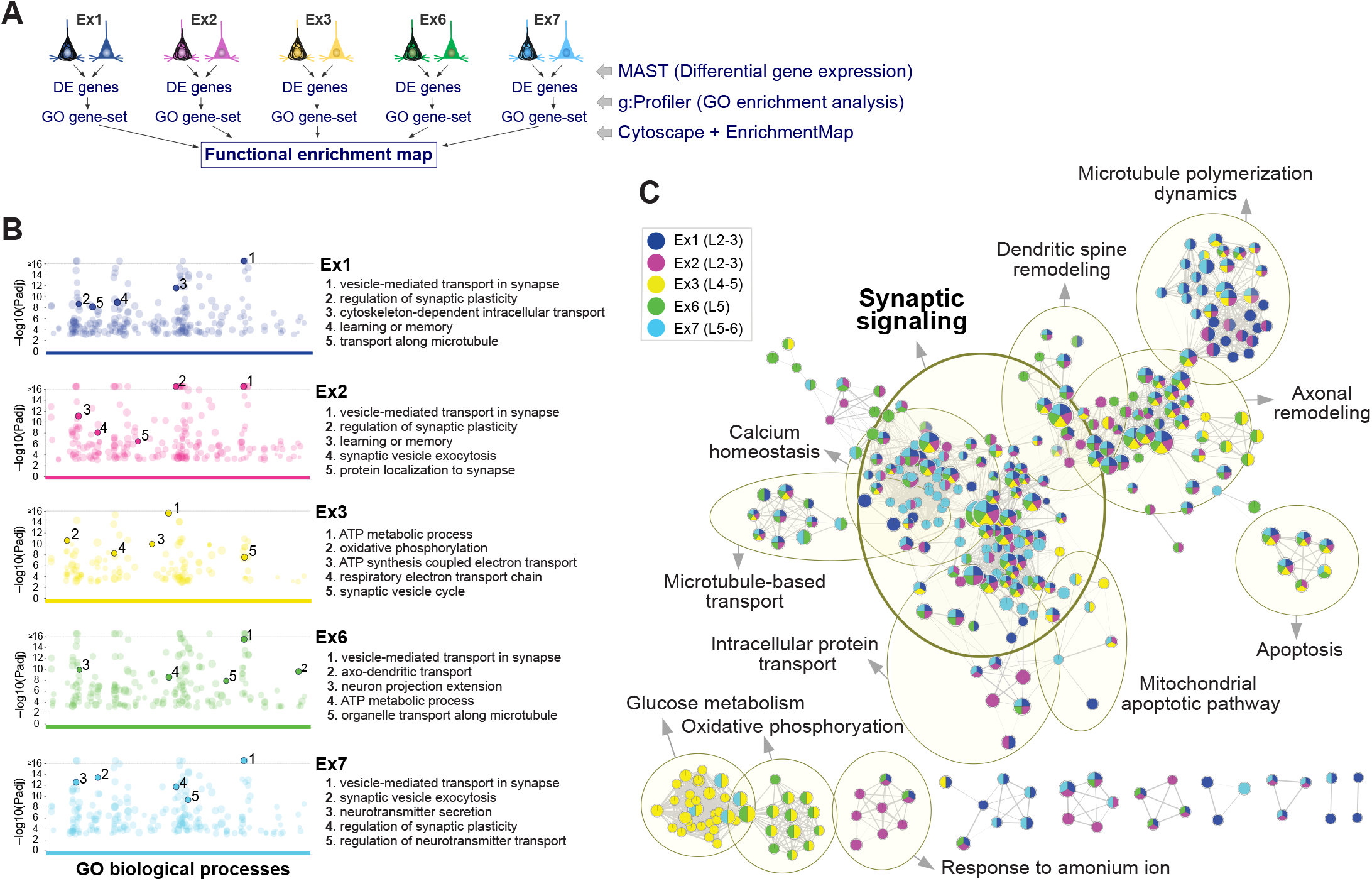
Shared versus cell-type-specific pathways associated with tau pathology. (A) Overview of our strategy to identify the shared and cell-type-specific pathways dysregulated in neurons with NFTs in the five excitatory clusters with the highest cell numbers and NFT proportions (E1, Ex2, Ex3, Ex6, Ex7). (B) Manhattan plots depicting the GO biological processes enriched in NFT compared to NFT-free neurons for each cell type obtained by g:Profiler. The y-axis shows the adjusted enrichment p-values on the −log10 scale and the x-axis represents functional terms. Colored circles represent significant terms (thresholds: g:SCS significance < 0.001; GO terms with > 50 or < 500 genes; capped at terms with a −log10 [Padj] > 16; Table S6). The top 5 nonredundant terms are highlighted. (C) Functional enrichment map generated from GO-derived gene sets using Cytoscape with the EnrichmentMap app. The input gene sets were the GO biological processes enriched in neurons with NFTs compared to those in NFT-free neurons within each cluster. Nodes represent the gene sets; the size of each node corresponds to the number of genes in the gene set (thresholds: p-value < 0.02; FDR q-value < 0.1). Edges between nodes represent overlapping genes between two gene sets; edge thickness represents the degree of overlap (overlap coefficient threshold of 0.7). Each cluster is color coded to illustrate the shared and distinct contributions of genes from each neuronal subtype. Yellow circles delineate constellations of functionally related gene sets, identified by the Autoannotate app. Constellations with > 5 nodes are represented. See also Tables S6 and S7.

### Dysregulation of synaptic transmission pathways in neurons with NFTs

To further characterize the transcriptome changes associated with synaptic transmission in neurons with NFTs, we used SynGO, a comprehensive, evidence-based reference for synaptic gene annotations and ontologies (Koopmans et al., 2019). Applying high-stringency parameters to filter annotations by experimental evidence, we identified 24 cellular component terms (171 genes) and 37 biological process terms (176 genes) significantly enriched at 1% false discovery rate (FDR) (Table S8). By cellular location, the postsynaptic density (PSD) membrane, presynaptic membrane, and presynaptic active zone were particularly overrepresented. The top-level overrepresented biological process terms were synapse organization, process in the presynapse and process in the postsynapse, whereas metabolism and transport were underrepresented (Figures 6A and S7). Specifically, the highest enrichment scores corresponded to the synaptic vesicle cycle (i.e., synaptic vesicle exocytosis and endocytosis), followed by transsynaptic signaling, synapse assembly, structural constituent of synapse, regulation of postsynaptic membrane receptor levels, and regulation of postsynaptic membrane potential (Figures 6A and 6B).

**Figure 6.**
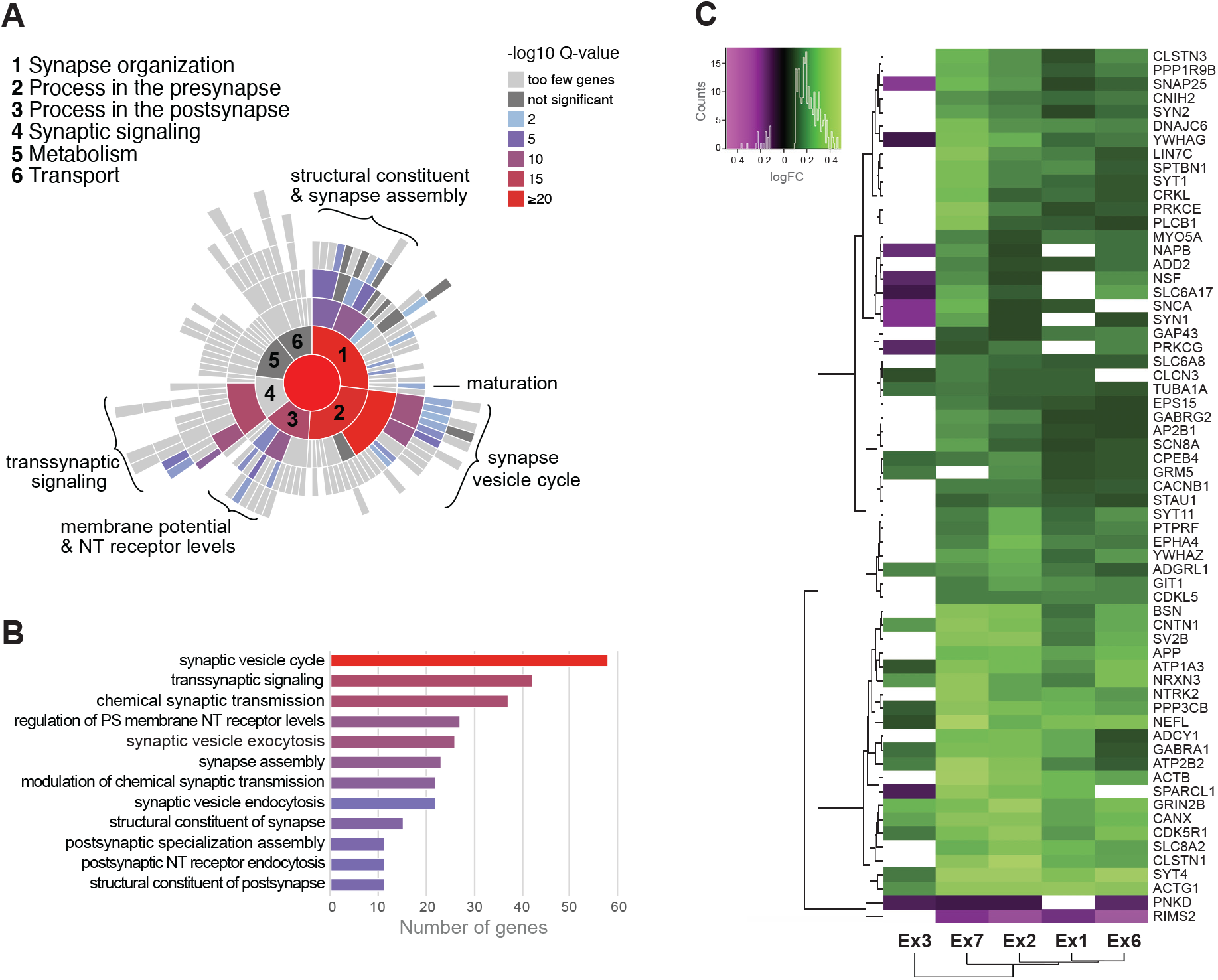
Dysregulation of synaptic transmission pathways in neurons with NFTs. (A and B) Enrichment of synaptic transmission pathways identified using SynGO annotations (Koopmans et al., 2019). The sunburst plot illustrates the over- and underrepresented biological process terms; colors represent enrichment values at 1% FDR. The bar plot shows the number of dysregulated genes and the enrichment values for the top 12 enriched terms. (C) Heatmap and hierarchical clustering of the core set of 63 synaptic genes dysregulated across five excitatory neuronal subtypes with high numbers of NFTs. Columns represent cell types; rows represent genes. The genes that are differentially expressed in ≥ 4 of the 5 clusters (63 genes) are represented (green = upregulated; magenta = downregulated; MAST test with adjusted p-value < 0.05; log fold change > 0.1; detection in ≥ 20% of cells). See also Figure S7 and Table S8.

A total of 227 DE genes in NFT neurons mapped to SynGO annotated genes (Table S8). The vast majority were upregulated (~89–95% in Ex1, Ex2, Ex6, and Ex7), except in Ex3 (~40% upregulated). Of the 227 DE genes, 17 were dysregulated in all 5 clusters, and an additional 46 were dysregulated in 4 of the 5 clusters (MAST test; adjusted p-value < 0.05; log fold change > 0.1; detection in ≥ 20% of cells) (Figures 6C and S7). Notably, among the commonly upregulated genes were genes encoding proteins that are biomarkers for neurodegeneration and cognitive decline in AD: NEFL, encoding neurofilament light chain; SNAP25, encoding synaptosomal‐associated protein 25, a SNARE complex protein with a role in synaptic vesicle exocytosis; and SYT1, encoding synaptotagmin-1, an integral membrane protein of synaptic vesicles implicated in calcium-dependent vesicular trafficking and exocytosis (Brinkmalm et al., 2014; Davidsson et al., 1996; Galasko et al., 2019; Mattsson et al., 2017; Mattsson et al., 2019; Preische et al., 2019) (Figures 6B and S7; Table S8). Other upregulated genes related to the synaptic vesicle cycle were the synaptotagmin genes STY4 and STY11; SV2B, encoding synaptic vesicle glycoprotein 2B; and BSN, encoding the presynaptic cytomatrix protein bassoon. A number of ion channels and membrane potential regulators were also commonly upregulated in NFT neurons, including genes encoding NMDA glutamate receptor subunits (GRIN2B and GRIN2A), GABAA receptor subunits (GABRA1 and GABRG2), Na^+^/K^+^–ATPase subunits (ATP1A3 and ATP2B2), and the voltage-gated sodium channel Nav1.6 (SCN8A). Other commonly upregulated genes included NTRK2, encoding neurotrophic receptor tyrosine kinase 2 (TrkB), a regulator of transsynaptic signaling activated by BDNF, and NRXN3, encoding the transsynaptic adhesion protein Neurexin 3. There were no genes downregulated in all 5 clusters and only 2 genes were downregulated in 4 of the 5 clusters: RIMS2, encoding a Rab3-interacting protein located in the presynaptic active zone and implicated in synaptic vesicle exocytosis, and PNKD, encoding a protein that interacts with RIM proteins and suppresses synaptic vesicle exocytosis (Kaeser et al., 2012; Shen et al., 2015) (Figures 6C and S7). Thus, our analysis identified a set of commonly dysregulated synaptic genes and pathways in neurons with NFTs that included well-established AD biomarkers, highlighting the value of our datasets for the discovery of novel biomarkers and therapeutic targets.

## Discussion

Despite the evidence linking tau pathology to histologic and neuroimaging features of neurodegeneration and cognitive decline in AD (Bejanin et al., 2017; Braak and Del Tredici, 2015; Franzmeier et al., 2019; Hanseeuw et al., 2019; Jack et al., 2018; Nelson et al., 2012; Schwarz et al., 2016; Schöll et al., 2016), the identity of the neurons that are susceptible to the development of tau pathology and their molecular alterations have not been resolved. To address this, we developed procedures for the isolation and transcriptome profiling of individual neurons with NFTs from human AD brain and profiled 63,110 NFT-bearing or neighboring NFT-free somas from the prefrontal cortex of eight Braak VI AD patients. Our study revealed a marked specificity for the neuronal subtypes and molecular pathways associated with tau aggregation in advanced stages of dementia. DGE analysis comparing NFT-bearing and NFT-free somas on a cell type basis identified the shared and cell-type-specific molecular signatures associated with NFTs. The top dysregulated pathways in NFT-bearing neurons were related to synaptic transmission, supporting the view of AD as a disorder of the synapse (de Calignon et al., 2010; de Wilde et al., 2016; Selkoe, 2002; Sheng et al., 2012; Spires-Jones and Hyman, 2014). Our datasets provide a framework for improved disease modeling and a resource for future explorations in basic and translational medicine related to tau pathology.

Our method allows for the unbiased, high-throughput isolation of neurons with NFTs from archived fresh-frozen brains. A caveat of our method is the disruption of cell membranes by freeze-thawing which results in the loss of most transcripts from the cytoplasmic compartment. Accordingly, our analysis comparing nuclear and soma FACS-sorted populations showed only modest increases in total UMIs and exonic reads (~16% and ~9.2%, respectively) in somas compared with nuclei, although cytoplasmic transcripts are estimated to comprise 50–80% of total transcripts (Bakken et al., 2018; Grindberg et al., 2013). The loss of cytoplasmic transcripts did not undermine the ability to discriminate between closely related neuronal subtypes, as previously shown in studies directly comparing the transcriptomes of nuclei and whole cells (Bakken et al., 2018; Lake et al., 2017). Variable amounts of cytoplasm, however, may introduce a bias in transcript quantification. To reduce technical variability, we compared NFT somas with neighboring NFT-free somas that were isolated and processed in parallel from the same tissue sample. This strategy also reduced potential confounding effects derived from patients’ genetic background, sex, age, comorbidities, medication, and premortem agonal state. Thus, we provide a new methodological assay to isolate and profile individual cells with cytoplasmic protein aggregates. Our method can be applied to other neurodegenerative tauopathies featuring neuronal and/or glial tau aggregates, such as progressive supranuclear palsy, corticobasal degeneration, Pick disease, primary age-related tauopathy, and chronic traumatic encephalopathy, to investigate shared and disease-specific mechanisms of tau-mediated neurodegeneration.

Previous studies have suggested several morphological and molecular features in cortical neurons that may underlie their different vulnerabilities to tau pathology. NFT formation has been associated with larger cell size, low expression of Ca^2+^-binding proteins, sparse myelinization, dysregulation of autophagy, and dysregulation of microtubule dynamics, among other effects (Fu et al., 2019; Mrdjen et al., 2019; Roussarie et al., 2018). Our study emphasized the specificity of distinct neuronal subtypes in the formation of NFTs. The subtypes with the highest NFT proportions included putative cortico-cortical projection neurons in layers 2-3 (clusters Ex1 and Ex2; CUX2^+^) and intratelencephalic projection neurons in layer 5 (cluster Ex6; RORB^+^/PCP4^+^) (Harris et al., 2019; Harris and Shepherd, 2015; Zeng et al., 2012). Among layer 6 cortico-thalamic neurons, deep layer 6b (Ex11; FEZF2^+^/CTGF^+^), a population that projects to anterior and mediodorsal (i.e., association) thalamic nuclei (Hoerder-Suabedissen et al., 2018; Zeng et al., 2012) in rodents, was relatively vulnerable. These results, together with previous neuropathology and functional imaging studies showing the stereotypical distribution pattern of tau pathology and its spread through functionally connected brain regions (Braak and Braak, 1991; Braak and Del Tredici, 2018; Franzmeier et al., 2019), support the association between tau pathology and neural connectivity. As previously suggested, vulnerable circuits may correspond to the default-mode network, a network of highly interconnected brain regions, including the limbic system, associative cerebral cortex, anterior and mediodorsal thalamic nuclei, and basal forebrain cholinergic system, which is highly active during the processing of different high-level cognitive tasks such as autobiographical episodic memory, social cognition, semantic processing and attention and is disrupted in AD (Alves et al., 2019; Buckner et al., 2008; Greicius et al., 2004; Raichle et al., 2001; Seeley et al., 2009).

Our DGE and pathway enrichment analyses highlighted the dysregulation of synaptic transmission, particularly the synaptic vesicle cycle, in NFT-bearing neurons across subtypes, supporting the previously reported association between tau pathology and synaptic function (Busche et al., 2019; Spires-Jones and Hyman, 2014; Wu et al., 2016). The central role of synaptic vesicle cycle dysregulation in the pathogenesis of AD has been suggested by other omic and cellular studies (Canchi et al., 2019; de Wilde et al., 2016; Zhou et al., 2017). Notably, we identified 227 synaptic genes dysregulated in neurons with NFT, including genes with key roles in the synaptic vesicle cycle (i.e., SNAP25, SV2B, SYT1, STY4, RIMS2, PNKD) and neuronal excitability (i.e., GRIN2A, GRIN2B, GABRA1, GABRG2, ATP1A3, ATP2B2, SCN8A). Correspondingly, we identified the TF regulatory network controlled by REST, a key regulator of neuronal excitability (Lu et al., 2014; Zullo et al., 2019), enriched in excitatory neurons with NFTs. Interestingly, more than ~90% of the DE genes in neurons with NFTs were upregulated, except in cluster Ex3 (~40% upregulated). Previous histological, protein and gene expression studies in AD postmortem tissue have consistently shown a loss of synapses and downregulation of synaptic markers during disease progression (Berchtold et al., 2013; Canchi et al., 2019; de Wilde et al., 2016; Grubman et al., 2019; Miller et al., 2017; Terry et al., 1991; Wang et al., 2016). Thus, the upregulation of synaptic genes identified in neurons with NFTs compared to NFT-free neurons may be at least in part the result of compensatory mechanisms in response to synaptic dysfunction and loss.

Whether tau aggregation is a driver of neurodegeneration or part of a protective response that ultimately fails to lead to neuronal death is an unsolved question. We identified the upregulation of the TF ATF4, the main effector of the integrated stress response, an adaptive pathway to restore cellular homeostasis (Pakos-Zebrucka et al., 2016). Other upregulated genes involved in the cellular stress response included APP, JUN, and HSP90AA1 and other genes encoding heat shock proteins (Klein et al., 2019; Mathys et al., 2019). As expected in association with protein misfolding, we identified both proapoptotic and negative regulators of apoptosis in neurons with NFTs. Although their ultimate effect in neuronal survival is to be determined, the overall enrichment of neuronal cell death and apoptosis pathways in NFT-bearing neurons was modest, consistent with the lack of acute death of neurons with aggregates observed in tau transgenic mice by *in vivo* multiphoton imaging (de Calignon et al., 2010).

In contrast to the shared dysregulation of synaptic and stress response genes in NFT-bearing neurons, we found that gene expression changes associated with metabolism and mitochondrial function were highly cell type dependent and particularly enriched in cluster Ex3. This cluster differed in the directionality of gene expression changes (mostly downregulated), the biological processes that were overrepresented (i.e, ATP metabolic process, oxidative phosphorylation, respiratory electron transport chain) and its TF regulatory networks (i.e, GABPA). Intriguingly, this cluster corresponds to a poorly characterized but distinct neuronal subtype in middle cortical layers that expresses RORB and PLCH1 and contains neurons expressing the gene encoding the neuropeptide galanin (GAL) (Alexandris et al., 2019).

Together, the results of our study revealed a marked selectivity of neuronal subtypes associated with tau pathology and common molecular pathways converging on the synapse. Although the pathogenic cascade leading to synaptic failure in AD is incompletely understood, APP cleavage products, Aβ oligomers, and tau oligomers have been shown to play key roles (Busche et al., 2019; Moore et al., 2015; Puzzo et al., 2017; Zott et al., 2019). We found an upregulation of APP in NFT-bearing neurons across excitatory subtypes. APP/Aβ is upstream of tau pathology in AD pathogenesis, and duplication of the APP gene is sufficient to cause early-onset AD (Franzmeier et al., 2019; Hardy and Selkoe, 2002; Moore et al., 2015; Rovelet-Lecrux et al., 2006). Thus, the gene expression changes identified in NFT-bearing neurons compared to neighboring NFT-free neurons may result from APP-dependent and tau-dependent mechanisms interacting at the single-cell level. NFT-bearing and NFT-free neurons may also respond differently to extracellular, soluble Aβ depending on their cell surface receptors. We identified the upregulation of potential receptors for Aβ in NFT-bearing neurons: PRNP, GRIN2B, GRIN2A, ATP1A3, EPHA4, and PGRMC1 (Fu et al., 2014; Izzo et al., 2014; Laurén et al., 2009; Ohnishi et al., 2015; Shankar et al., 2007). Together, our findings support the idea that interactions between APP and tau pathways at the single-cell level lead to synaptic dysfunction and neurodegeneration. Although we cannot distinguish drivers of degeneration from compensatory responses, our transcriptome data provide a resource for exploring pathogenic mechanisms and a platform for biomarker and drug discovery.

## Acknowledgments

Human brain tissue was obtained from the UCLA-Easton Center and the NIH Neurobiobank (Sepulveda repository, Los Angeles, CA). We thank Abel Nunez, Spencer Tung, and Kazu Williams for their kind assistance providing the tissue blocks. FACS was performed at the UCLA Jonsson Comprehensive Cancer Center (JCCC) and Center for AIDS Research Flow Cytometry Core Facility. 10x Genomics Chromium and sequencing was performed at the UCLA Technology Center for Genomics & Bioinformatics (TCGB).

## Author contributions

M.O.G. and I.C. designed all experiments; M.O.G. and I.C. developed the methods to isolate and profile somas with tau aggregates; M.O.G. and I.C. processed samples for single-soma RNA-seq with assistance of Y.D.; M.O.G., I.C., T.A., W.E.L., and R.K. performed the computational analysis; S.M. and V.S. performed the TF enrichment analyses; Y.X. performed the validation studies in human brain tissue with assistance of T.S.; I.C. supervised the project and wrote the manuscript with input from all co-authors.

## Data Availability

The sequencing raw data and the digital expression matrices obtained using the 10x Genomics software Cell Ranger are available in the NCBI’s Gene Expression Omnibus (GSE129308) and are accessible through GEO Series accession number GSM3704357-GSM3704375. The computer scripts to reproduce the analysis and figures are available from the authors on reasonable request.

## Methods

### Isolation of individual somas with tau aggregates from frozen human brains

Fresh-frozen brain tissue blocks stored at –80°C were first warmed to –12°C to enable the dissection of thick (~500 µm) tissue sections while preserving the remaining frozen tissue for additional experiments. For each experiment, a section of the cortex (~200 mg) encompassing an equal representation of all cortical layers was cut. The tissue was dissected under a stereomicroscope to remove the white matter and leptomeninges and was then chopped into small pieces (< 1 mm^3^) using a chilled razor blade. To prevent further RNA degradation, all steps were performed on ice in RNase-free conditions. For tissue homogenization, a Potter-Elvehjem tissue grinder was used. These grinders have a clearance space between the pestle and tubes (0.1-0.15 mm clearance; 8 mL tubes) wider than those of the grinders that are typically used to dissociate nuclei, which facilitated the dissociation of relatively well-preserved somas. Each tissue sample was dissociated using 2.4 mL of homogenization buffer (10 nM Tris pH 8, 5 mM MgCl2, 25 mM KCl, 250 mM sucrose, 1 μM DTT, 0.5x protease inhibitor [cOmplete, Roche #4693159001], and 0.2 U/μL RNase inhibitor). No enzymatic digestion or detergents were used. For this amount of tissue, ~15 grinder strokes were needed. The number of strokes was adjusted by microscopically assessing the number and morphology of somas and the presence of clumps using a hemocytometer. Homogenates were then filtered through a 100-μm cell strainer and transferred into two 1.5-mL Eppendorf tubes.

Further clean-up was performed using iodixanol gradient centrifugation. The homogenate was first centrifuged at 400 ×g for 5 min at 4°C, and then, the supernatant was aspirated and discarded, and the pellets were gently resuspended in 200 μL of cold homogenization buffer. The homogenates were pooled into one tube, and the total volume was measured and adjusted with homogenization buffer to obtain an exact volume of 450 μL. An equal volume (450 μL) of 42% v/v iodixanol medium (75 mM sucrose, 25 mM KCl, 5 mM MgCl2, 10 mM Tris [pH 8], and 42% w/v iodixanol) was added to the homogenate and gently mixed with a pipette to obtain a final concentration of 21% iodixanol. The mixture was then transferred to a new 2-mL Eppendorf tube containing 900 μL of cold 25% iodixanol medium (146 mM sucrose, 48 mM KCl, 10 mM MgCl2, 19 mM Tris [pH 8], and 25% w/v iodixanol) by slow layering on the top. The tubes were centrifuged at 8,000 ×g for 15 min at 4°C, resulting in the sedimentation of somas at the bottom, covered by the supernatant and a top layer of thick material containing cell clumps and abundant myelin. The top layer and supernatant were removed and discarded carefully, avoiding contamination of the pellet. Pellets were detached with a small amount (~50 μL) of immunostaining buffer (0.1 M phosphate-buffered saline [PBS; pH 7.4], 0.5% bovine serum albumin [BSA], 5 mM MgCl2, 2 U/mL DNAse I, and 0.2 U/μL RNase inhibitor), transferred to clean tubes, and gently resuspended in a total volume of 200 μL of immunostaining buffer. After a 15-min incubation with immunostaining buffer, at 4°C, with gentle rocking, primary antibodies were added (mouse anti-phospho-Tau [Ser202, Thr205] monoclonal antibody [AT8], 1:150, ThermoFisher cat#MN1020; rabbit anti-MAP2 polyclonal antibody, 1:40, Millipore cat #AB5622), and the suspension was incubated for 40 min at 4°C with gentle rocking. An equal volume (500 μL) of immunostaining buffer was then added, and the tubes were inverted several times before being centrifuged at 400 ×g for 5 min at 4°C. The supernatant was carefully removed, and the pellets were resuspended in 600 μL of immunostaining buffer. Secondary antibodies (goat-anti-mouse, Alexa Fluor 350, 1:500; goat-anti-rabbit, Alexa Fluor 647, 1:500) and a nuclear stain (SYTOX green, 1:40,000) were added, and the solutions were incubated for 30 min at 4°C with gentle rocking. Aliquots of unstained, only secondary antibody-treated, and single-stained (SYTOX green, MAP2, or AT8) cells were saved for use as FACS controls.

The number and morphology of the somas were evaluated microscopically after each critical step and immediately before FACS. Good-quality samples contained a suspension of single somas and naked nuclei, with few clumps and little debris; the proportion of cells with relatively well-preserved somas varied between 20-50% of the total sample. The typical yield for ~100 mg of cerebral cortex tissue was between 0.5–1.5 × 10^6^ somas.

### FACS of somas with and without NFTs

FACS was used to collect single-cell suspensions of somas with tau aggregates (AT8^+^) and neighboring neurons without tau aggregates (MAP2^+^/AT8^−^). Sorting was performed using a BD FACSAria II at the Flow Cytometry Core Laboratory at UCLA. PBS was used as the sheath fluid, with a sheath pressure of 20 psi. A 100-μm nozzle tip was used, and the frequency of droplet generation was ~30 kHz. The primary laser was a blue Trigon 488-nm used in the generation of forward scatter (FSC) and side scatter (SSC). The secondary lasers were UV Trigon 355-nm, blue Trigon 488-nm, and red Trigon 640-nm, used for the excitation of Alexa 350, SYTOX green, and Alexa Fluor 647, respectively. Sample events were acquired at < 30% droplet occupancy.

FACS gates were based on a combination of regions drawn around target populations in 2D plots, performed in the following order: FSC height vs. SSC height; SSC area vs. FSC width; FSC area vs. FSC width; SSC area vs. SYTOX green fluorescence (bandpass filter 525/50); and Alexa 350 fluorescence (bandpass filter 450/50) vs. Alexa Fluor 647 fluorescence (bandpass filter 670/30). The FSC versus SSC gates were set with permissive limits, discarding the smallest and largest particles. SYTOX green fluorescence was used to discriminate single cells from cell clumps and anucleated cell fragments. Alexa Fluor 647 was used to discriminate neurons with soma (MAP2^+^) from nonneuronal cells and naked neuronal nuclei (MAP2^−^). Alexa Fluor 350 was used to discriminate between somas containing tau aggregates (AT8^+^) and somas without tau aggregates (AT8^−^). Unstained, only secondary antibody-treated, and only single primary antibody-treated cell suspensions were included as controls to minimize false positives due to nonspecific staining or autofluorescence.

Two populations were collected: AT8^+^ (either positive or negative for MAP2) and MAP2^+^/AT8^−^ somas. The yield per sample ranged from 1,600–37,000 somas for AT8^+^ and over 3 × 10^5^ somas for MAP2^+^. Somas were collected in 1.5-mL Eppendorf tubes containing 100–200 μL of collection buffer (0.1 M PBS [pH 7.4] and 0.1 U/μL RNase inhibitor). After collection, BSA was added to each tube for at a final concentration of 1%. To prevent somas from adhering to the tube walls, the Eppendorf tubes used for collection were precoated with BSA by filling tubes with 10% BSA solution in PBS for 5 min, rinsing with PBS, and drying at 4°C overnight.

### Single-soma RNA-sequencing and data analysis

Single-soma RNA capture and library preparation were performed using the 10x Genomics Chromium Single Cell 3’ v2 assay. Single-cell suspensions for FACS were centrifuged at 400 ×g for 5 min at 4°C to concentrate cells. Without disturbing the pellet, sufficient supernatant was removed to achieve a concentration of ~350 cells per μL. Cell concentrations were measured using a hemocytometer, and the quality of cells was examined under a fluorescence microscope. The numbers of loaded cells ranged from 1,400–11,000 to capture the maximum number of cells, with an upper limit of ~5,000 cells per sample (for an expected cell capture efficiency of ~40%). The following steps were performed according to the manufacturer’s instructions. For cDNA amplification, the number of PCR cycles used was 13–15 (adjusted to the targeted cell recovery). For library construction, the number of cycles for the sample index PCR was 12–13 (adjusted for the quantified cDNA input).

The generated paired-end libraries were sequenced on an Illumina Novaseq 6000 using two lanes of an S2 flow cell (2 × 100). We sequenced 26 bp for read 1 (cell barcode and UMI), 8 bp for the i7 index, and 98 bp for read 2. All libraries were combined and sequenced together in a single run; the concentration of each sample was normalized to the total number of cells to achieve similar numbers of reads per cell. Cells were sequenced at a depth of ~72,000 reads per cell, corresponding to a sequencing saturation of ~85%.

Paired-end sequence reads were processed using the 10x Genomics software package Cell Ranger version 3.0.1. We used the Cell Ranger count pipeline with default parameters to perform alignment to the prebuilt reference genome GRCh38 and for filtering, barcode counting, and UMI counting (see Table S2 for sequencing quality control metrics). The resulting digital expression matrices were analyzed using the R-based Seurat package, version v2.3.

To analyze the MAP2^+^ datasets, we loaded all digital expression matrices into Seurat, filtered out cells with < 250 genes or > 12,000 UMIs, and removed mitochondrial DNA-encoded genes, ribosomal genes, and uncharacterized RP11-, RP13-, RP1-, RP3-, RP4-, RP5-, and RP6-genes. All datasets were combined and normalized using the function LogNormalize with the default scale factor 10,000. The number of UMIs and samples of origin were regressed out. We selected genes with average expression values between 0.0075 and 3 and dispersion values > 0.3 (~4,200 genes) for downstream analysis. Principle component analysis (PCA) was used to reduce dimensionality, and the first 22 statistically significant principal components (PCs) were selected for clustering. Clusters were identified using a graph-based clustering approach (Seurat FindClusters function with the following parameters: 1:22 PCs; 1.0 resolution; 100 random start positions and 10 iterations per random start; and 30 k for the k-nearest neighbor algorithm) and visualized with t-SNE using the same PCs. Cluster-specific marker genes were obtained by comparing the gene expression levels for each individual cluster with those for all other cells using the Wilcoxon rank-sum test. Genes detected in ≥ 25% of cells (in either the tested cluster or in all other cells combined) with positive log fold changes > 0.25 and adjusted p-values < 0.05 were included. Cluster robustness was assessed by examining cluster stability after subsetting and rerunning clustering and by comparing our data with previous human brain single-nucleus RNA-seq (Hodge et al., 2019; Mathys et al., 2019) and gene expression data. We annotated 21 clusters corresponding to 11 excitatory neuron subtypes, 7 inhibitory neuron subtypes, and 3 glial cell types. Clusters containing cells with mixed identities and/or cell states (1.57%; gray colored) were removed from further analysis.

To analyze the AT8^+^ datasets and the combined MAP2^+^ and AT8^+^ datasets, we used multiCCA for dimensionality reduction (Butler et al., 2018). MultiCCA performed better that PCA in distinguishing between cell types and disease states. We used the same parameters as described above to filter low-quality cells, except that we regressed out mitochondrial DNA-encoded genes instead of filtering them. The top 4,200 highly variable genes that were present in at least two datasets were selected for downstream analysis. The first 26 canonical correlation vectors were aligned using the datasets/samples as a grouping variable (Figures S4G–J). Clustering and t-SNE were performed using the same canonical correlation and a resolution of 1. Cluster-specific marker gene identification and cluster robustness assessment were performed as described above.

To further assess the robustness of the clustering of the combined MAP2^+^ and AT8^+^ datasets, we reanalyzed our sequencing data after alignment to a pre-mRNA reference genome (Figure S4). This approach has been shown to markedly improve gene detection in nucleus preparations due to the high fraction of intronic reads captured in nuclear sequencing (Bakken et al., 2018). We created a custom pre-mRNA reference package by modifying the prebuilt reference genome GRCh38 provided by Cell Ranger to include both intronic and exonic reads for downstream analysis. The resulting digital expression matrices were analyzed using version v3.1 of the R-based Seurat package. We used the single-cell transform (SCT) method for normalization, variance stabilization, regression of the number of UMIs and mitochondrial gene fraction (Hafemeister and Satija, 2019), and uniform manifold approximation and projection (UMAP) for dimensionality reduction and clustering. By selecting the first 6 statistically significant PCs, we first obtained three clusters corresponding to excitatory neurons, inhibitory neurons and nonneuronal cells. Subsequently, we performed an independent clustering analysis for the excitatory neuron and inhibitory neuron subsets. Identification of cluster-specific marker genes and cluster annotation was performed as described above (Figures S4B–D).

### DGE analysis

DGE between cells with and without NFTs within each cell type was assessed using the MAST generalized linear model (Finak et al., 2015) on each cluster separately (Table S4). MAST takes into account the characteristic bimodal distribution of single-cell data in which gene expression is either detected (nonzero) or not detected (typically high due to the high rate of dropout events) and has been shown to perform highly favorably on statistical power and FDR control compared to other methods for DGE analysis in single-cell datasets (Soneson and Robinson, 2018). We applied the following cut-off values to generate the lists of DE genes: adjusted p-value < 0.05; log fold change (positive or negative) > 0.1; and detection in ≥ 20% of cells for at least one condition. To visualize shared and distinct DE genes among cell types, we generated gene expression heatmaps by hierarchical clustering of genes using Ward’s minimum variance method with the heatmap.2 R package. The resulting clustering was used to build the row (genes) dendrogram. Columns (cell types) were clustered using Euclidian distance and reordered by mean values.

Cross-validation of the molecular signatures associated with NFTs was performed by rerunning DGE after excluding a subset of samples. We removed different combinations of two of the eight samples from the dataset, generated full lists of DE genes using MAST, and compared them with the lists of DE genes generated from the 8-sample dataset. We used the RRHO (Plaisier et al., 2010) to identify and visualize statistically significant overlap between pairs of gene lists (RRHO 1.22.0 R package). The full lists of genes without any cut-off filters were ranked by their adjusted p-values and the statistical significance of the number of overlapping genes measured successively to determine the strength and pattern of correlations. The output was visualized using heatmaps with a step size (i.e, resolution) of 50 (Figure S5).

### TFBS analysis

Transcription factor binding site (TFBS) enrichment analysis was performed in the promoters of the genes that were DE in cells with NFTs, compared to cells that were NFT free in five neuronal subtypes (Ex1, Ex2, Ex3, Ex6, and Ex7), using the TFBS pipeline described elsewhere (Parikshak et al., 2016). The region 1 kb upstream of the transcription start site was defined as the canonical promoter region. For each of the five clusters, we assessed the top 200 connected genes, ranked by intramodular connectivity (kME), using the Religious Orders Study and Memory and Aging Project (ROSMAP) prefrontal cortex AD dataset (Mostafavi et al., 2018). Putative motifs bound by the TF were obtained from the TRANSFAC database (Matys et al., 2003). We identified the upstream sequences of these 200 genes using the Clover algorithm (Frith et al., 2004) to calculate motif enrichment. The background for enrichment was calculated with the MEME algorithm (Bailey and Elkan, 1994) using 1000-bp sequences upstream of all human genes, human CpG islands, and the sequence of human chromosome 20. We calculated p-values by selecting 1,000 sequences of the same length, testing them for enrichment using MEME, and computing the p-values based on the observed motif enrichment ranks versus the randomized sets. The enriched TFs for each of the five neuronal subtype clusters were obtained (Figure S6A).

### TF regulatory networks

To generate neuronal TF regulatory networks, we obtained data for human neuronal TFs and their target genes from the ENCODE ChIP-seq dataset (Consortium, 2012; Davis et al., 2018) and intersected them with the TFs identified in our TFBS enrichment analysis. For each TF regulatory network, we defined the genes with robust evidence of coexpression across brain tissues based on AMP-AD network analysis using the top 100 connected genes (ranked by kME) for each coexpression module (Morabito, 2019) the genes that were DE in cells with NFTs versus NFT-free cells (Table S4), and cluster-specific background genes (expressed in ≥ 10% of cells in the cluster). The edges in the networks represent bicorrelation of gene expression values in the AMP-AD datasets, and the nodes are spatially arranged by multidimensional scaling (MDS) of the AMP-AD gene expression. In the interest of visual clarity of the networks, we limited the genes displayed to a maximum of 25 DE genes (ranked by p-value) and 20 background genes (ranked by p-value); the numbers of DE genes between NFT versus NFT-free cells that were also coexpressed in the AMP-AD network analysis were not limited (Figure S6B).

### GO enrichment analysis

The web server g:Profiler (version Ensembl 97, Ensembl Genomes 44) (Raudvere et al., 2019) was used to perform GO enrichment analysis. For each cluster, we input the list of DE genes between cells with and without NFTs against a list of background genes (expressed in ≥ 10% of cells in the cluster) and obtained hierarchical sorting lists of GO terms. Statistical significance thresholds were determined using Fisher’s exact test and multiple testing correction (default native method g:SCS). To limit the size of functional categories subjected to enrichment analysis, we filtered out GO terms with < 50 or > 500 genes. Data were downloaded in generic enrichment map (GEM) format (Table S6) to be used as input for functional enrichment analysis.

Cytoscape with EnrichmentMap (Merico et al., 2010) was used to integrate and visualize GO enrichment results from the five excitatory clusters with the highest cell numbers and percentages of AT8^+^ cells into a single network. The ranked lists of statistically significant GO biological process terms obtained with g:Profiler were loaded into Cytoscape v7.3.1 with the EnrichmentMap app. v3.2 using the following conservative parameters: nodes (representing GO-derived gene sets) included gene sets with p-values < 0.02 and FDR q-values < 0.1; edges (representing gene overlap between gene sets) used an overlap coefficient threshold of 0.7. Each of the five clusters was color coded to visualize the shared and distinct contributions of each cell type. The enrichment map was annotated automatically using the Autoannotate app. and the clusters labeled with three words using the WordCould app.

The SynGO enrichment tool (Koopmans et al., 2019) was used to further characterize the synaptic transmission pathways enriched in neurons with NFTs. The list of genes associated with synaptic transmission by functional enrichment analysis (genes in clusters #1, #2 and #6 in Table S7; 510 genes) was loaded against a custom background list containing all genes expressed in the five excitatory cell subtypes analyzed (6,843 genes; expressed in ≥ 10% of cells). Using stringent settings and a minimum of 3 matching input genes per term, 227 genes mapped to SynGO annotated categories; 171 genes had a cellular component annotation, and 176 genes had a biological process annotation. A total of 24 cellular component and 39 biological process terms were significantly enriched at 1% FDR.

### Histological validation in human brain tissue

Validation of gene expression was performed in prefrontal cortex tissue from AD patients using RNAscope ISH. Single and double chromogenic ISH staining was performed on 20-μm-thick cryosections of fresh-frozen prefrontal cortex tissue following the manufacturer’s protocol (RNAscope 2.5 HD assay and duplex assay). Human RNAscope probes were obtained from ACD Bio. to detect the following genes: SLC17A7 (#415611), GAD1 (#404031), CUX2 (#425581), LAMP5 (#487691), COL5A2 (#510911), PCP4 (#446111), ROBO3 (#483191), NR4A2 (#582621), and NTNG2 (custom designed to target nucleotides 1217-2210 of NM_032536.3 with 20 Z pairs). Adjacent Nissl-stained and NeuN-immunostained sections served as anatomical references to delineate boundaries between cortical layers.

Quantification of the proportions of neurons with NFTs in specific subtypes was performed by dual or triple fluorescent AT8 immunohistochemistry and RNAscope ISH for cell identity markers (SLC17A7, GAD1, CUX2, PCP4, ROBO3, NR4A2, NTNG2) on 12-μm-thick sections from four AD patients. Prior to staining, the sections were photobleached to quench lipofuscin autofluorescence using an LED light source (Sun et al., 2017). For this, sections were fixed in 4% paraformaldehyde for 15 min at 4°C, washed twice in 1x PBS for 5 min, and then exposed to an LED light source (300-watt, full spectrum LED; Platinum LED Lights, cat# P300). Sections were kept in 1x PBS, placed at a distance of 40 cm from the LED, and exposed for 36–48 hours at 4°C. After photobleaching, RNAscope ISH was performed using the Multiplex Fluorescent Reagent Kit v2 according to the manufacturer’s instructions, except for a shortened protease treatment time of 15 min. Fluorescence signals were amplified and visualized using the TSA Plus Cyanine-5 and the TSA Plus Fluorescein systems (Akoya Biosciences, #NEL745E001KT and #NEL741E001KT, respectively), according to the manufacturer’s instructions, using a TSA Plus working solution concentration of 1:500. After ISH, the sections were fixed in 4% paraformaldehyde for 15 min at 4°C and then washed twice in 1x PBS for 5 min. Nonspecific binding was blocked with 10% normal goat serum in PBS for 30 min at 4°C. Sections were then incubated with 1:100 AT8 antibody in 1x PBS with 5% normal goat serum at 4°C overnight. The next day, the sections were washed three times for 10 min in 1x PBS and incubated with 1:50 goat anti-mouse Alexa Fluor 350 for 1 hour at 4°C. Sections were washed in 1x PBS three times for 5 min and mounted with Vectashield antifade mounting medium (Vector Laboratories).

Quantification was performed on digital images taken at 400x magnification with a Zeiss Axio Imager M2 microscope equipped with a monochrome digital camera (Hamamatsu C11440-22CU) and the Zeiss ApoTome.2 optical sectioning system. To quantify the colocalization of AT8 with different neuronal subtype markers, multiple images were acquired automatically within a region of interest that was traced manually to include the entire thickness of the cortex (for SLC17A7 and GAD1), the upper layers 2-3 (for CUX2), the middle layers 3b-5 (for PCP4), layer 5b (for PCP4 and ROBO3), or layer 6 (for NTNG2 and NR4A2) and then combined into a single image using the slide-scanning module in Stereo Investigator software v.2018 (MBF Bioscience). A total of 10–12 counting frames (400 µm x 250 µm) were randomly placed within the region of interest to cover an area of ~5 mm^2^. Double- or triple-positive cells were counted manually using the Placing markers module in Stereo Investigator. For PCP4/ROBO3/AT8 triple staining, cells were counted throughout layer 5b (7.5 to 15 mm2). Four patients and ≥ 2 sections per marker were analyzed. The investigator was blinded to the sample and to the results obtained from single-cell RNA-seq studies. A total of 1,200–5,000 cells per marker were analyzed. The results are expressed as the percentage of double- or triple-positive cells and the standard deviation for each gene marker.

## Supplemental information

**Figure S1.**
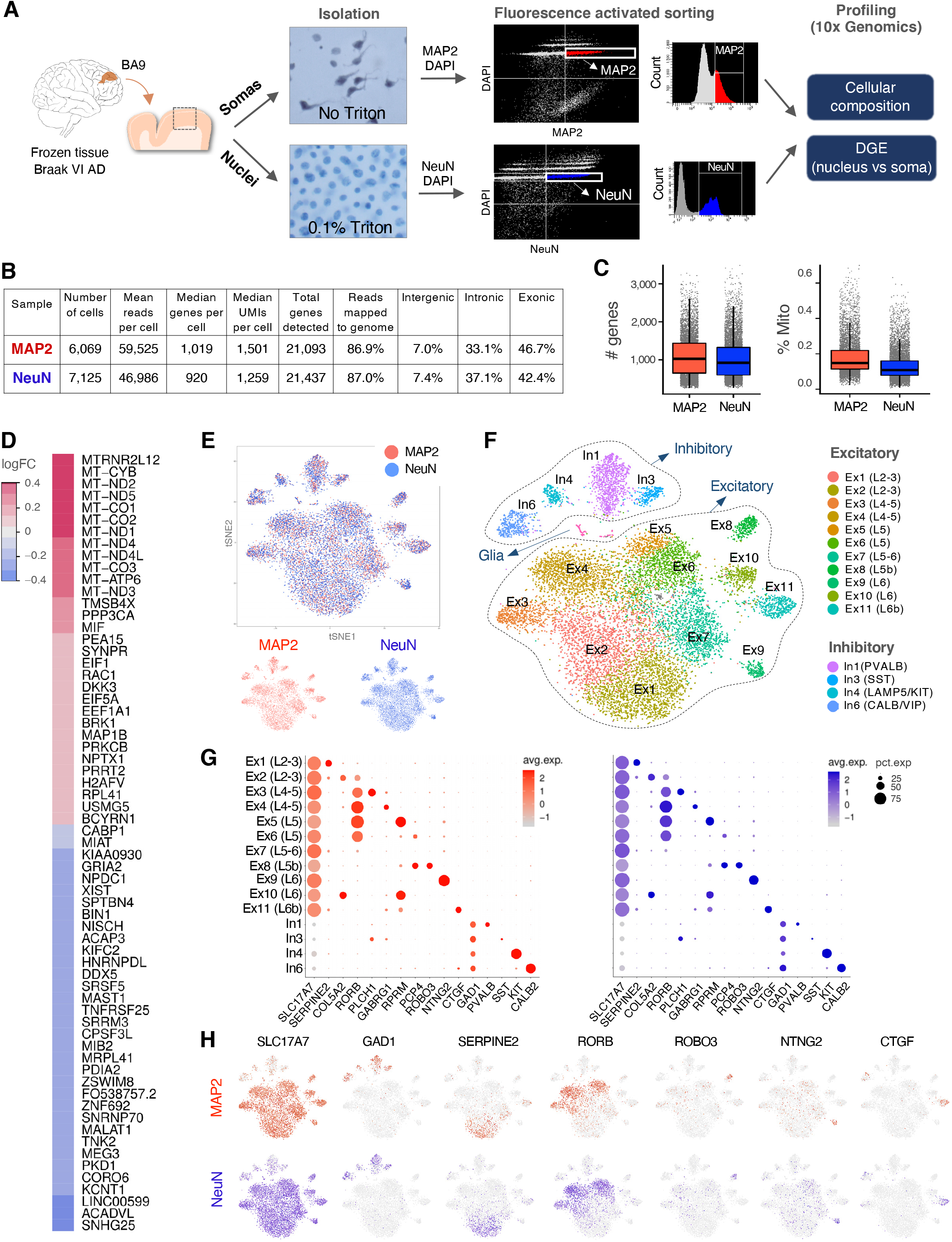
Comparison between NeuN^+^ (nuclei) and MAP2^+^ (somas) transcriptomes. (A) Overview of the experimental approach used to compare the transcriptomes from single nuclei (NeuN^+^) and single somas (MAP2^+^) from the BA9 of an AD Braak stage VI subject (case # 6). Nuclei were dissociated using mechanical force (Dounce tissue grinder; clearance space between pestle and tight tube: 0.02–0.056 mm) in the presence of 0.1% Triton X-100; somas were dissociated with mechanical force only (Potter-Elvehjem tissue grinder; clearance space between pestle and tube: 0.1–0.15 mm). Microphotographs illustrate single-nucleus and single-soma suspensions. Nuclei were stained with DAPI and immunostained with NeuN (nuclear); somas were stained with DAPI and immunostained with MAP2 (cytoplasmic). The two populations were isolated by FACS, and their transcriptomes were profiled using 10x Genomics Chromium Single Cell 3’ v2. (B) Table showing the estimated numbers of nuclei and somas and the sequencing metrics obtained with the Cell Ranger pipeline (10x Genomics). (C) Box plots representing the numbers and distributions of genes and mitochondrial transcript fractions in each sample. Each dot represents a single cell. (D) Heatmap showing the DE genes between MAP2 and NeuN datasets (red: overrepresented in MAP2; blue: overrepresented in NeuN; Wilcoxon rank test with Bonferroni-corrected p-value ≤ 0.05, average log fold change ≥ 0.1; detection in ≥ 10% of cells for at least one condition). (E) t-SNE plots combining the two samples (coded by color), showing similar clustering patterns between the nucleus and soma datasets. (F) Unbiased clustering identified the same cell types (11 excitatory neuron clusters [Ex1 to Ex11; 11,051 cells], 4 inhibitory neuron clusters [1,895 cells], and a few glial cells [109 cells]). (G) Dot plots from MAP2 (left) and NeuN (right) datasets depicting highly similar expression levels of several marker genes (x-axis) within major excitatory (SLC17A7^+^) and inhibitory (GAD1^+^) neuronal subtypes (y-axis). The sizes of the dots represent the percentage of neurons expressing the marker; color intensities represent scaled expression levels. (H) t-SNE plots showing similar expression patterns of SLC17A7, GAD1, SERPINE2, RORB, ROBO3, NTNG2, and CTGF in the MAP2 (top) and NeuN (bottom) datasets.

**Figure S2.**
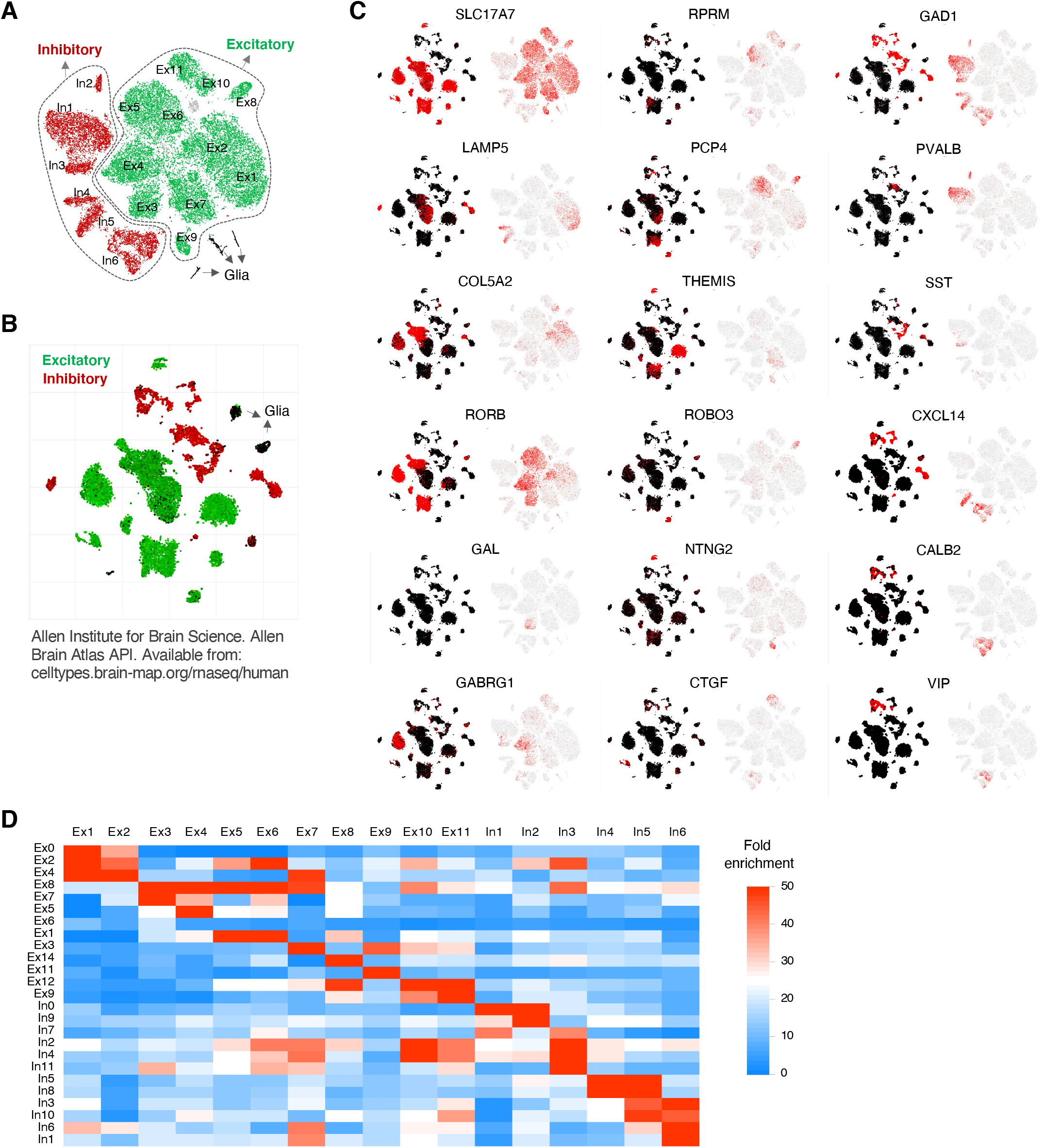
Cluster annotations compared to reference datasets. (A and B) t-SNE plots showing unbiased clustering of excitatory and inhibitory neuronal subtypes in our MAP2 dataset (A; clusters as in Figure 2C; 38,465 somas from eight brains; age 57-93 years; BA9; sequenced with 10x Genomics) compared to a reference dataset from healthy controls (B; 15,928 nuclei from eight brains; age 24-66 years; middle temporal gyrus; sequenced with SMART-Seq) (Hodge et al., 2019). (C) t-SNE plots showing comparable expression patterns of excitatory and inhibitory marker genes in both datasets. (D) Heatmap comparing neuronal subtype annotations in our MAP2 dataset (columns) with those from a previously reported AD dataset (rows) (Mathys et al., 2019).

**Figure S3.**
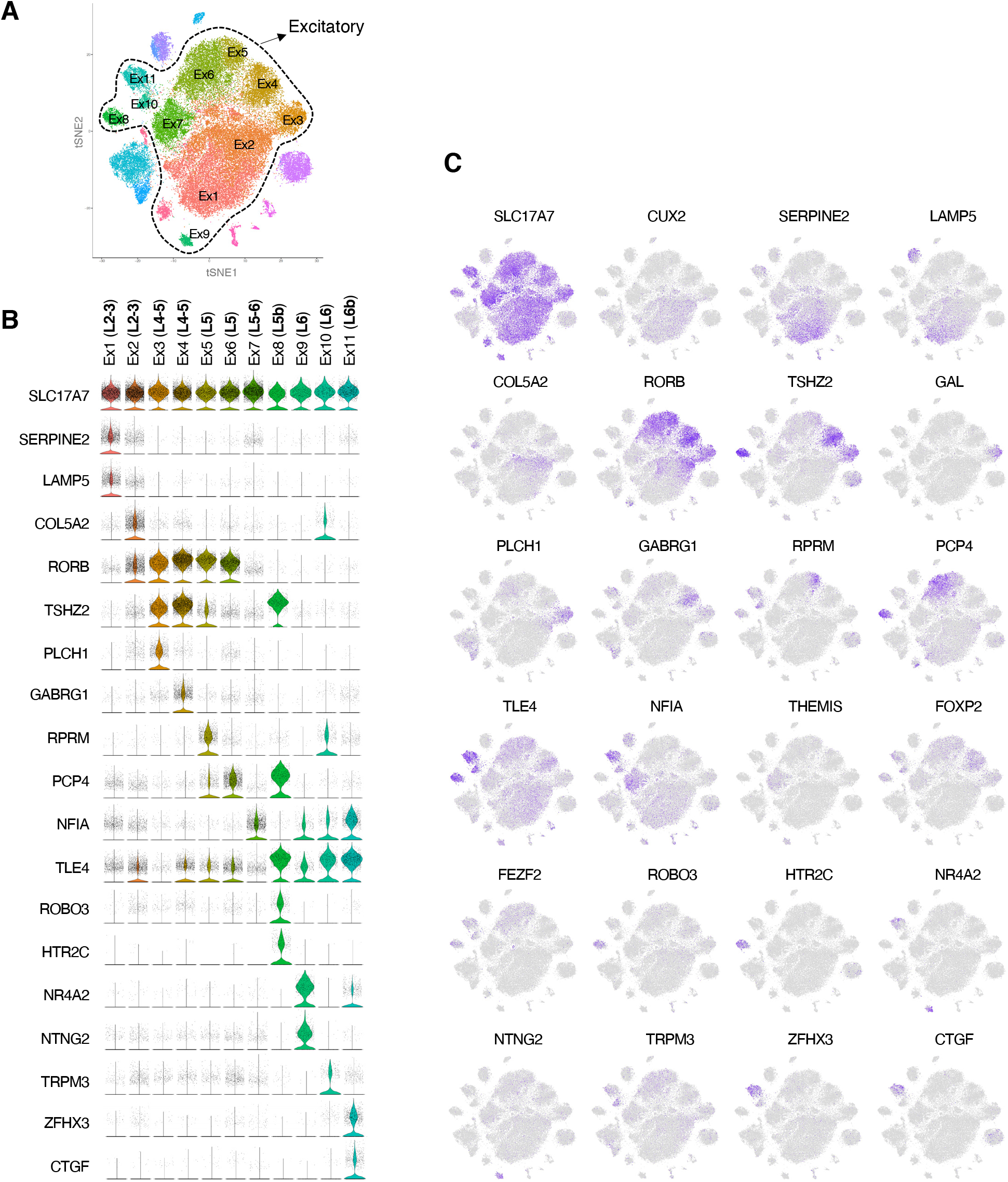
Gene markers for excitatory cell clusters. (A) t-SNE plot showing 11 excitatory neuron clusters (Ex1-Ex11), identified as in Figure 2I. (B and C) Violin plots and t-SNE plots showing the expression of 24 marker genes for excitatory neuronal subtypes.

**Figure S4.**
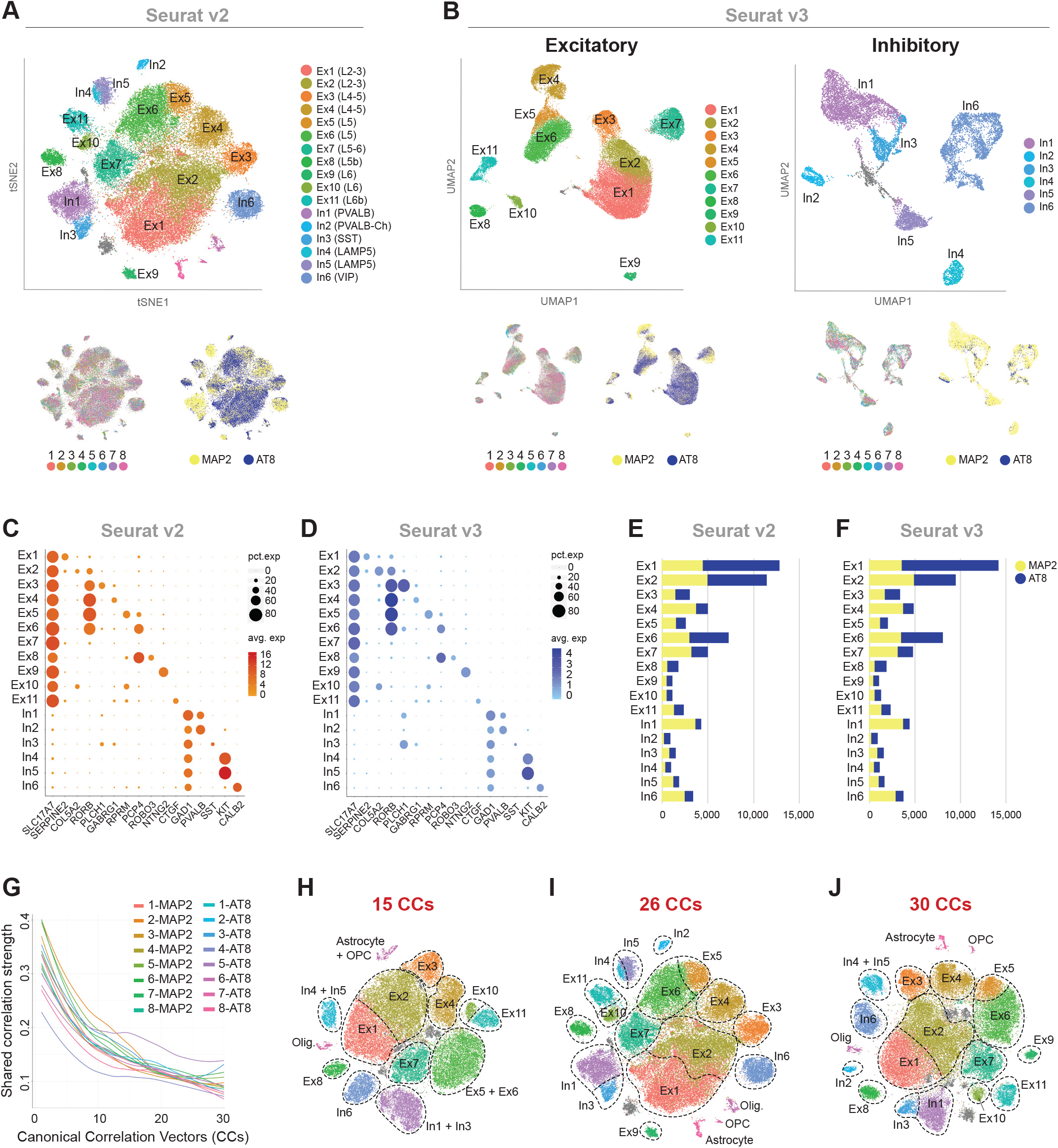
Robustness of clustering of the NFT and NFT-free combined dataset. (A–F) Comparison of unsupervised clustering using two strategies. The NFT and NFT-free combined dataset was analyzed using either exonic sequences for alignment and multiCCA for data integration and clustering (A, C, and E; Seurat v2.4 package) or pre-mRNA for alignment and the Seurat v3 workflow with SCTransform for data integration and clustering and uniform manifold approximation and projection (UMAP) for dimensionality reduction and clustering (B, D, and F; Seurat v3.1 package). Colors in the t-SNE plots in (A) correspond to cell types (top), subjects (bottom left) or datasets (bottom right). The same color code is used in (B), where the excitatory clusters (left) and inhibitory (right) clusters were subset and processed separately. Clusters were annotated as in Figure 2C (11 excitatory neuron subtypes and 7 inhibitory neuron subtypes) in both workflows. The dot plots in (C and D) depict similar expression of marker genes (x-axis) within each excitatory and inhibitory neuronal subtype (y-axis) from the combined dataset processed with either Seurat v2 (left) or Seurat v3 (right). The sizes of the dots represent the percentage of neurons expressing the marker; color intensities represent expression levels. The bar plots in (E and F) show the frequencies of neurons with NFTs within each cluster in the datasets processed with either Seurat v2 (left) or Seurat v3 (right). Bars represent the absolute numbers of MAP2^+^ (yellow) and AT8^+^ (blue) somas for each excitatory and inhibitory neuronal subtype. Clustering robustness is shown by the similar results in cell type identification and NFT frequencies obtained with both workflows. (G–J) Selection of canonical correlation vectors (CCs) for dimensionality reduction in the multiCCA analysis of the NFT and NFT-free combined dataset. The plot in (G) represents the correlation strength between samples as a function of the number of CCs. Color corresponds to samples (MAP2 and AT8 datasets; 8 samples each; 63,110 somas in total). This visualization method is used to select the number of CCs for dimensionality reduction. The t-SNE plots demonstrate clustering after selecting 15 (H), 26 (I) or 30 (J) CCs. Color corresponds to cell types. Clustering robustness is shown by the wide range of CCs producing similar clustering results. MultiCCA analysis with 26 CCs was chosen for downstream analysis.

**Figure S5.**
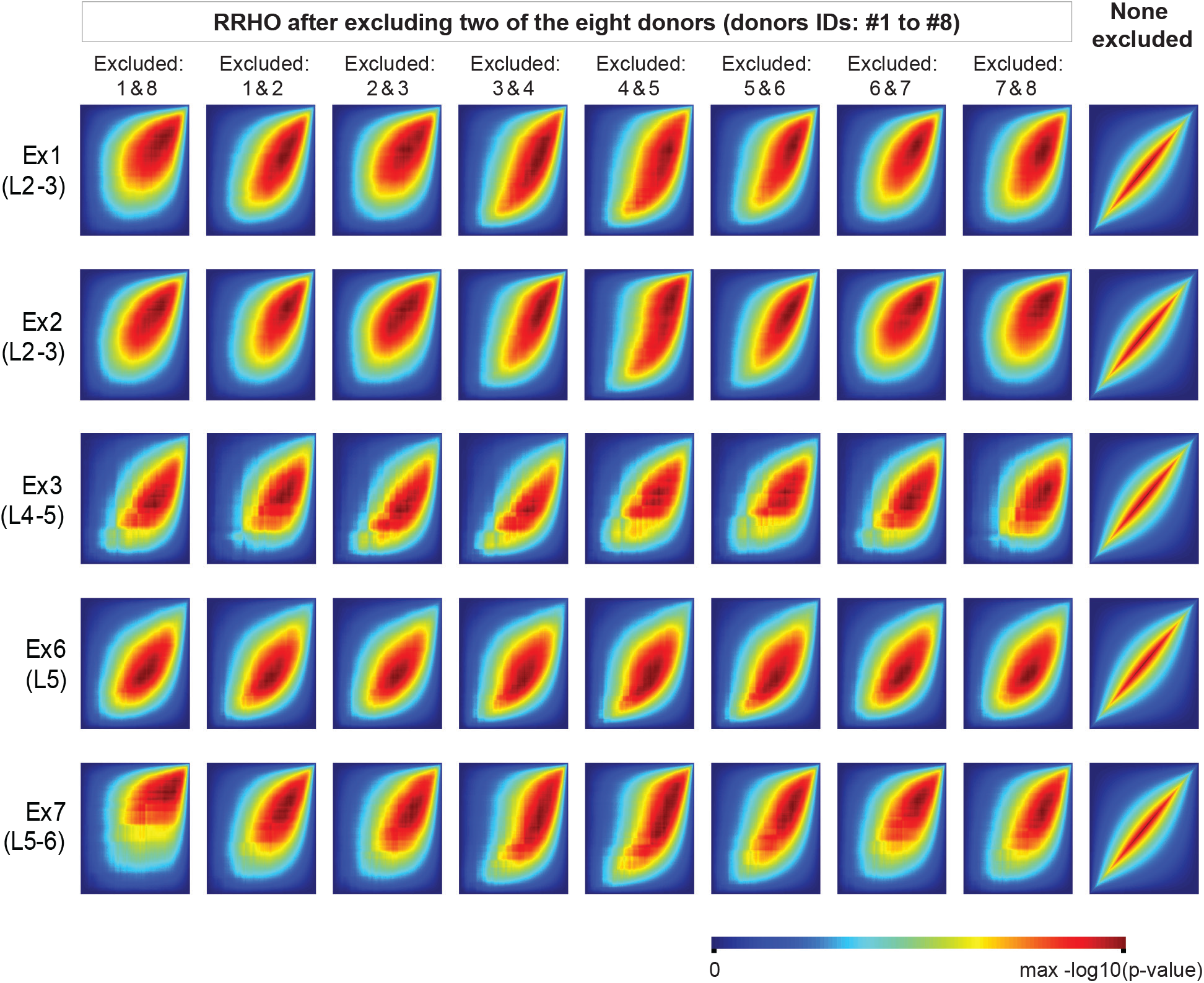
Assessment of the robustness of transcriptional changes associated with NFTs by subsampling. Rank-rank hypergeometric overlap (RRHO) heatmaps representing the overlap or correlation between the gene expression signatures associated with NFTs obtained from the dataset combining the eight donors (IDs #1 to #8) and the gene expression signatures obtained after excluding two donors (top x-axis = excluded donors). In the far-right column, no samples were excluded, and the heatmaps depict a perfect correlation. Rows correspond to the five excitatory clusters with a high burden of tau pathology (Ex1, Ex2, Ex3, Ex6, and Ex7). Gene expression signatures are robust to sub-sampling of any two of the eight donors.

**Figure S6.**
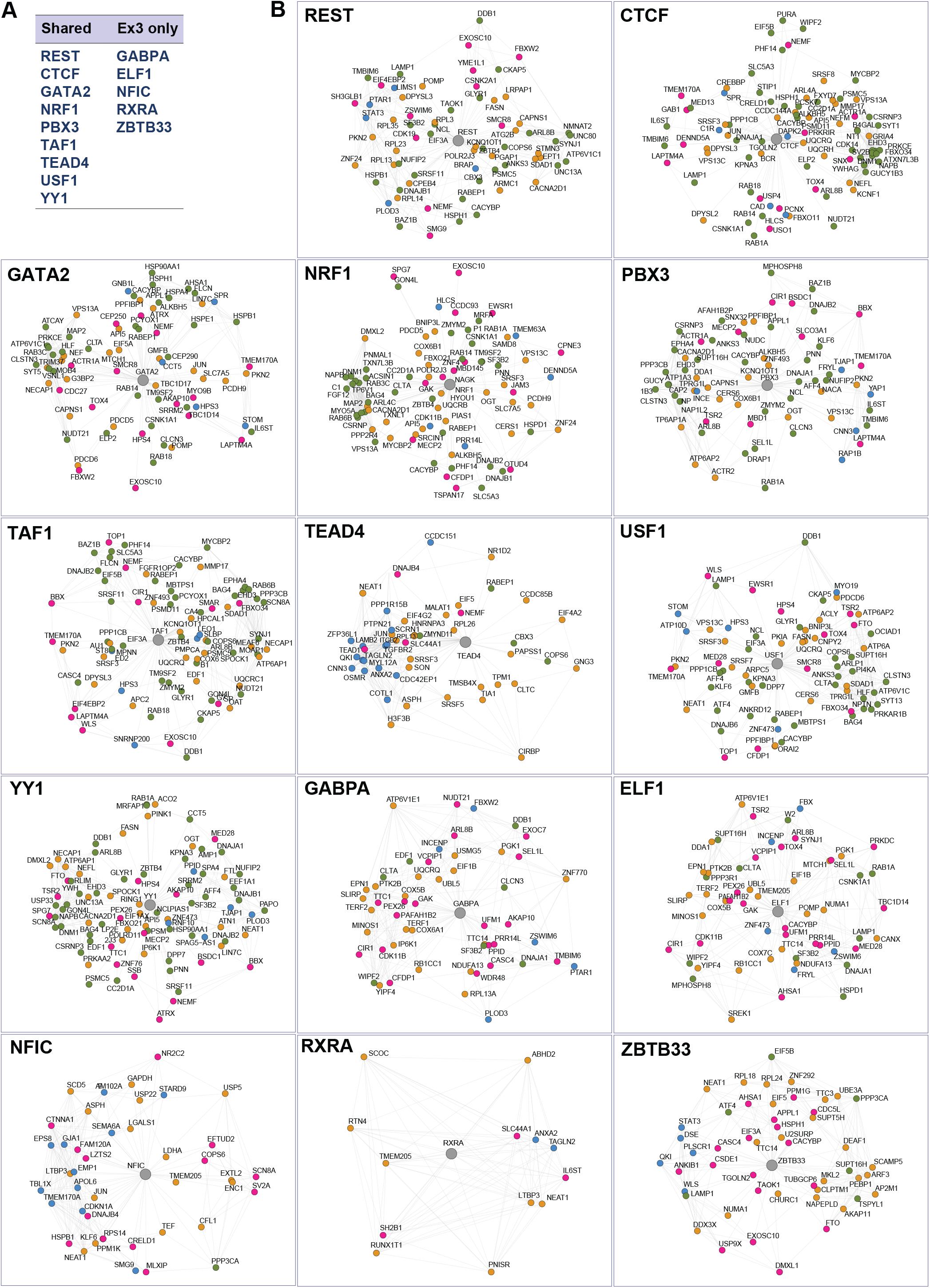
Transcriptional regulatory networks associated with NFTs. (A) Transcription factors identified by a TFBS enrichment analysis performed in the promoters of the genes that were differentially expressed in NFT-bearing compared to NFT-free neurons in five neuronal subtypes (Ex1, Ex2, Ex3, Ex6, and Ex7). The transcription factors enriched in the five excitatory clusters and those unique to cluster Ex3 are shown. (B) Transcriptional regulatory network plots (color code: yellow = coregulated and differentially expressed in NFT-bearing cells; pink = coregulated and coexpressed based on our single-cell data; green = coregulated, differentially expressed in NFT-bearing cells, and coexpressed based on ROSMAP datasets; blue = coregulated and coexpressed based on ROSMAP datasets).

**Figure S7.**
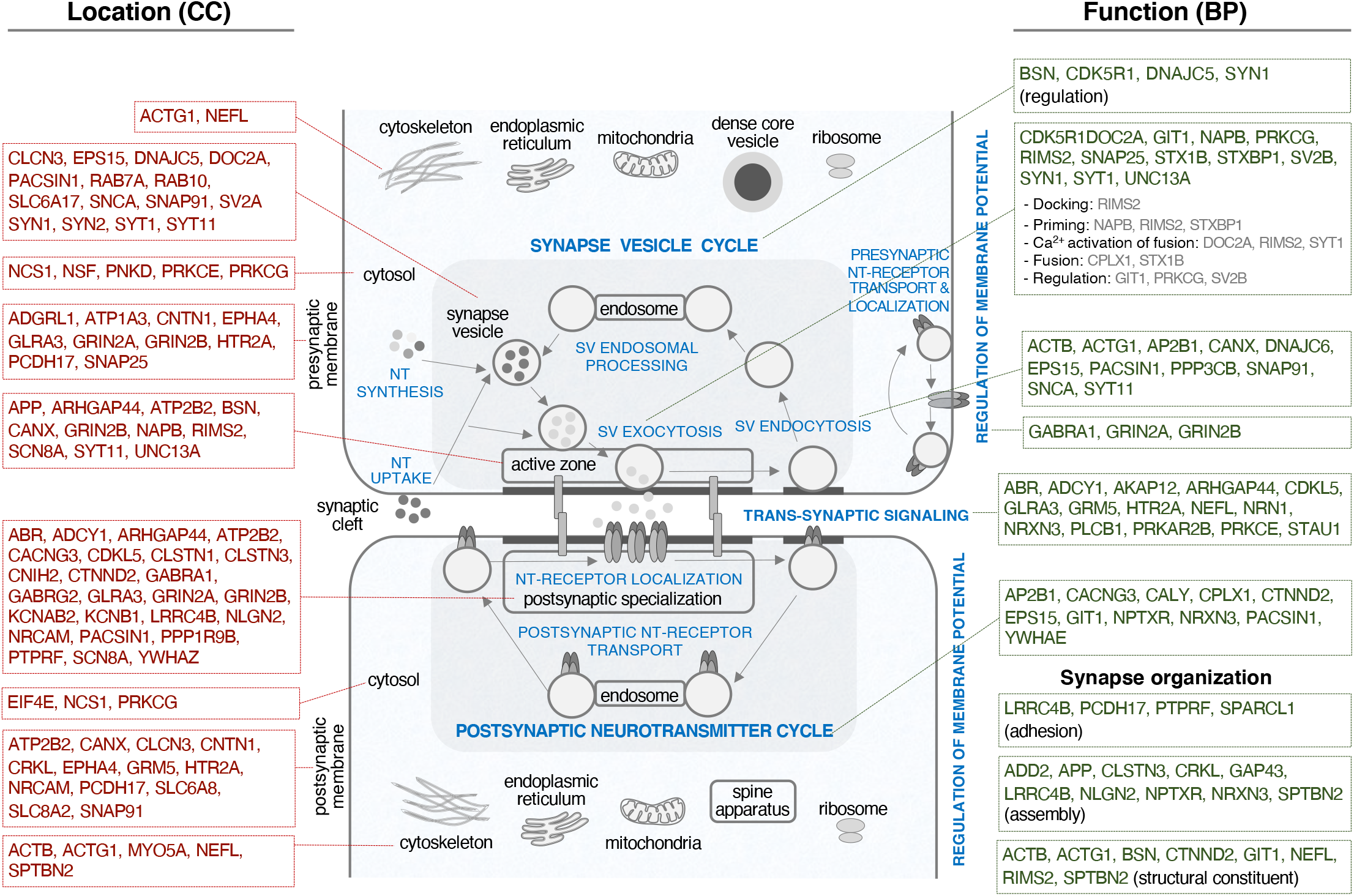
Mapping of synaptic genes dysregulated in NFT-bearing neurons on a model synapse. Schematic representation of a model synapse adapted from the SynGO framework (Koopmans et al., 2019). Top-level GO cellular component (CC) and biological process (BP) annotations are depicted. The DE genes between neurons with and without NFTs in five excitatory clusters with a high burden of tau pathology (Ex1, Ex2, Ex3, Ex6, and Ex7) that were significantly enriched in the synapse (SynGO analysis; data from Figure 6 and Table S7) are mapped onto their corresponding CC and/or BP. Only the genes that were differentially expressed in at least three of the five clusters are represented. Most genes were upregulated, except RIMS2, PNKD, and CALY. A few genes were downregulated in cluster Ex3 and upregulated in the other clusters (e.g., SNAP25, SLC6A17, YWHAG, SNCA, SPARCL1, SYN1, NAPB, NSF, PRKCG).

**Table S1. AD subject demographics**

Characteristics of the AD donors selected for the single-cell profiling of neurons with NFTs. All patients died with dementia and received a neuropathological diagnosis of Alzheimer disease neuropathological change, a Braak stage VI of VI, and an ABC score (NIA-AA Research Framework criteria) of A3B3C3 (Braak and Braak, 1991; Hyman et al., 2012). No neocortical Lewy body pathology was present. Brains #1 and #7 showed cardiovascular disease (CVD), including moderate to severe atherosclerosis in large vessels, moderate to severe arteriosclerosis, and rare remote lacunar infarcts. Cortical infarcts were not seen by gross examination, and the cerebral cortical tissue used for single-cell RNA-seq experiments was assessed histologically to ensure the absence of infarcts within or in close proximity to the sampled tissue. Brains #4, #6 and #8 were from subjects with early-onset dementia. The postmortem interval (PMI) ranged from 1 to 19.5 hours. The RNA integrity number (RIN) ranged from 5.7 to 7.0, as measured by an Agilent Bioanalyzer using the total RNA Nano kit. Abbreviations: M = male; F = female.

**Table S2. Single-cell sequencing metrics**

Estimated numbers of cells and sequencing and alignment metrics for each sample, obtained with the cellranger count pipeline from Cell Ranger (10x Genomics).

**Table S3. Gene markers and cell counts**

Clusters obtained from the MAP2^+^, AT8^+^, and combined AT8^+^/MAP2^+^ datasets. Gene markers include those genes detected in ≥ 25% of cells, with a positive log fold change > 0.25, and an adjusted p-value < 0.05. Cell counts show the numbers of cells contained in each cluster.

**Table S4. Differential gene expression between cells with and without NFTs**

DE genes between cells with and without NFTs within each cluster. DE genes were defined as those genes detected in ≥ 20% of cells in at least one condition, with a log fold change (positive or negative) > 0.1 and an adjusted p-value < 0.05, based on the MAST test. Background genes were defined as those detected in ≥ 10% of cells in at least one condition.

**Table S5. Interactive hierarchical heatmap of DE genes between cells with and without NFTs across cell types**

Heatmap of DE genes across neuronal subtypes (clusters Ex1, Ex2, Ex3, Ex4, Ex5, Ex6, Ex7, Ex8, and Ex11). Columns represent cell types; rows represent genes. Green = upregulated; magenta = downregulated. Genes are hierarchically clustered by log fold change values. Individual genes can be highlighted by clicking on the gene name on the right y-axis. Moving the cursor over each box shows the gene name (row), cell type (column), and log fold change value (value). DE genes were defined as those genes detected in ≥ 20% of cells in at least one condition, with a log fold change > 0.1 and an adjusted p-value < 0.05, based on the MAST test.

**Table S6. GO enrichment analysis comparing cells with and without NFTs**

GO terms from five excitatory clusters with a high burden of tau pathology (Ex1, Ex2, Ex3, Ex6, and Ex7). The lists of enriched GO biological process terms were generated with g:Profiler using as input the list of DE genes between cells with and without NFTs (genes expressed in ≥ 20% cells; log fold change > 0.1; adjusted p-value < 0.05; data from Table S4) against background genes (expressed in ≥ 10% of cells; data from Table S4). Columns show GO biological process terms (filtered to include terms containing 50–500 genes), genes, and p-values, using default g:SCS significance thresholds.

**Table S7. Functional enrichment analysis**

Genes, nodes and annotated clusters from an enrichment map integrating five excitatory cell types with a high burden of tau pathology (Ex1, Ex2, Ex3, Ex6, and Ex7). The enrichment map was generated using Cytoscape and EnrichmentMap. Nodes included gene sets with p-values < 0.02 and FDR q-values < 0.1; edges used an overlap coefficient threshold of 0.7. Clusters were annotated automatically using the Autoannotate app. Spreadsheets labeled cluster nodes show GO IDs, GO terms, number of genes, and gene names within each annotated cluster. Spreadsheets labeled cluster genes show heatmaps displaying gene expression across the five cell types (green = expressed; black = not detected).

**Table S8. Enrichment analysis of synaptic transmission genes in neurons with NFTs by SynGO**

Synaptic transmission genes and pathways enriched in NFT neurons compared to NFT-free neurons in five excitatory clusters with a high burden of tau pathology (Ex1, Ex2, Ex3, Ex6, and Ex7). This file contains the input and custom background gene lists for SynGO analysis (input list = 510 genes, dysregulated in neurons with NFTs as identified in the functional gene enrichment analysis [Table S7]; custom background list = 6,843 genes, expressed in >10% cells of any of the five excitatory neuronal subtypes), the output gene list from SynGO (227 genes mapped to SynGO annotated genes using stringent settings and a minimum of 3 matching input genes per term), and a ranked list of the DE genes between cells with and without NFTs identified by SynGO.

