## Supplementary material for "Single-soma transcriptomics of tangle-bearing neurons in Alzheimer’s disease reveals the signatures of tau-associated synaptic dysfunction": Table S1

| Subject ID # | Age | Sex | PMI (hr) | RIN | Brain weight | Brain region | AD stage (NIA-AA) | Braak stage | Other pathology |
|---|---|---|---|---|---|---|---|---|---|
| **1** | 93 | F | 4 | 6.5 | 1,150 | BA9 | A3B3C3 | VI | CVD |
| **2** | 79 | F | 19.5 | 7.0 | 1,270 | BA9 | A3B3C3 | VI | none |
| **3** | 81 | M | 16 | 6.5 | 1,360 | BA9 | A3B3C3 | VI | none |
| **4** | 57 | M | 14 | 6.8 | 1,160 | BA9 | A3B3C3 | VI | none |
| **5** | 81 | F | 24 | 6.1 | 960 | BA9 | A3B3C3 | VI | none |
| **6** | 73 | F | 13 | 5.7 | 1,300 | BA9 | A3B3C3 | VI | none |
| **7** | 89 | F | 1 | 6.5 | 1,130 | BA9 | A3B3C3 | VI | CVD |
| **8** | 62 | M | 11 | 6.6 | 1,250 | BA9 | A3B3C3 | VI | none |

**Table S1. AD subject demographics**

Characteristics of the AD donors selected for the single-cell profiling of neurons with NFTs. All patients died with dementia and received a neuropathological diagnosis of Alzheimer disease neuropathological change, a Braak stage VI of VI, and an ABC score (NIA-AA Research Framework criteria) of A3B3C3 (Braak and Braak 1991; Hyman et al. 2012). No neocortical Lewy body pathology was present. Brains #1 and #7 showed cardiovascular disease (CVD), including moderate to severe atherosclerosis in large vessels, moderate to severe arteriosclerosis, and rare remote lacunar infarcts. Cortical infarcts were not seen by gross examination, and the cerebral cortical tissue used for single-cell RNA-seq experiments was assessed histologically to ensure the absence of infarcts within or in close proximity to the sampled tissue. Brains #4, #6 and #8 were from subjects with early-onset dementia. The postmortem interval (PMI) ranged from 1 to 19.5 hours. The RNA integrity number (RIN) ranged from 5.7 to 7.0, as measured by an Agilent Bioanalyzer using the total RNA Nano kit. Abbreviations: M = male; F = female.
